# Multimodally trackable and clinically translatable platform for modelling human demyelinating brain diseases by temporally dispersed chemically induced lesions in the pig brain

**DOI:** 10.1101/2023.07.26.550644

**Authors:** Mihai Ancău, Goutam Kumar Tanti, Vicki Marie Butenschoen, Jens Gempt, Igor Yakushev, Stephan Nekolla, Mark Mühlau, Christian Scheunemann, Sebastian Heininger, Benjamin Löwe, Erik Löwe, Silke Baer, Johannes Fischer, Judith Reiser, Sai S. Ayachit, Friederike Liesche-Starnecker, Jürgen Schlegel, Kaspar Matiasek, Martina Schifferer, Jan S. Kirschke, Thomas Misgeld, Tim Lueth, Bernhard Hemmer

## Abstract

**Background:** Despite advances in therapy, inflammatory demyelinating diseases of the central nervous system, such as multiple sclerosis, remain important causes of morbidity among young adults. Translation of remyelinating paradigms from current murine models is encumbered by the small size and low white matter content of the brains, limiting the spatial resolution of diagnostic imaging. Large animal models might be more suited for this purpose but pose significant technological, ethical and logistical challenges.

**Method:** We induced reversible and targeted cerebral demyelinating lesions by controlled injection of lysophosphatidylcholine in the minipig brain. One strength of the approach is the serial induction, allowing parallel imaging of successive stages of de-/remyelination.

**Findings:** We demonstrate controlled, clinically unapparent, reversible and multimodally trackable brain white matter demyelination in a large animal model. Lesions were amenable to follow-up using the same clinical imaging modalities (3T magnetic resonance imaging, ^11^C-PIB positron emission tomography) and standard histopathology protocols as for human diagnostics, as well as electron microscopy to compare against biopsy data from two patients with cerebral demyelination.

**Interpretation:** By employing human diagnostic tools and validating the model against data from related human diseases, our platform overcomes one important translational barrier of current animal brain demyelination models while having the potential for developing diagnostic procedures and imaging biomarkers. Remyelination and axon preservation dynamics diverge from classical rodent models.

**Funding:** This work was supported by the DFG under Germany’s Excellence Strategy within the framework of the Munich Cluster for Systems Neurology (EXC 2145 SyNergy, ID 390857198) and TRR 274/1 2020, 408885537 (projects B03 and Z01).

**Research in context:** *Evidence before this study:* Inflammatory demyelinating diseases of the central nervous system (CNS), targeting primarily the white matter (WM) of the brain and spinal cord, such as multiple sclerosis (MS), still represent some of the most important non-traumatic causes of disability in young adults. Current animal models based on murine species, for example, experimental autoimmune encephalomyelitis, have been demonstrated to reliably depict pathophysiological facets of human disease. However, they are nevertheless encumbered by the low WM content and the small size of murine brains, which still pose a translational barrier to diagnostic imaging tools used in a clinical context in human patients. Minipigs are increasingly being used to model human neurological diseases, as yet primarily in the context of neurodegenerative disorders.

*Added value of this study:* Here, we establish a platform for Minipig Stereotactic White-matter Injection using Navigation by Electromagnetism (MiniSWINE) and validate such a tool in a clinical multimodal imaging and microscopy setting against biopsy and imaging data from human demyelinating disorders across different disease stages, as well as against existing and potentially emerging human diagnostic imaging. Moreover, in order to overcome the neuroanatomical challenges of stereotactic injection in the pig brain, we designed a new electromagnetic-guided tracking system whose key advantage is the direct measurement of the injection cannula tip position in situ. Another strength of our study lies in its setup, characterized by the serial induction of successive stages of de- and remyelination, allowing for multimodal assessment via imaging and histopathology or electron microscopy of multiple stages in parallel. The remyelination dynamics inferred in this context diverge from the classical rodent studies, by exhibiting incomplete remyelination at the subacute stage, persistent astroglial and microglial activation as well as a minor degree of secondary axonal degeneration. Thus, they more closely resemble human inflammatory demyelinating brain plaques.

*Implications of all the available evidence:* We believe that MiniSWINE links evidence from well-established demyelination-induction methods from rodent models of CNS demyelinating disorders, as well as from human imaging and biopsy data, while at the same time providing a novel platform for the potential development of diagnostic procedures, discovery of imaging biomarkers and testing of remyelinating agents in diseases such as MS. Thus, it can have particular relevance to human health in the context of future translational animal model-based research in inflammatory demyelinating disorders of the CNS. Additionally, our electromagnetic-guided injection technique may enhance stereotactic substance delivery in human neurosurgery.

## Introduction

Inflammatory demyelinating disorders of the central nervous system (CNS), including multiple sclerosis (MS), neuromyelitis optica (NMO), acute disseminated encephalomyelitis (ADEM) or MOG-antibody-associated disease (MOGAD), represent the most common non-traumatic causes of disability in young adults, afflicting more than 2.5 million people worldwide(1). Magnetic resonance imaging (MRI) allows for assessing lesions in the brain and spinal cord in these diseases and thus has become an essential tool for clinical care. However, it still is not fully understood which MRI signatures relate to which histopathological stage of lesion evolution, such as the key transition from demyelination to remyelination(2). Refining the imaging readout for remyelination in early clinical trials is one of the main challenges limiting the development of remyelinating therapies for MS(3). While rodent models of demyelinating CNS disorders would be logistically convenient to devise novel diagnostic MRI- and positron emission tomography (PET)-techniques, these models are not optimal for immediate translation because of the small brain size, relative lack of white matter (WM) and the lissencephalic nature of rodent brains, as well as due to genomic and proteomic discrepancies between rodents and humans(4–6).

These limitations could be overcome using a large animal model that is more similar to human neuroanatomy. Indeed, for translational progress, a model would be optimal in which successive stages of lesion de- and remyelination could be detected with the same diagnostic protocols and, ideally, devices as in human clinical studies. Here, we report such a model based on refined stereotactic microinjection of lysophosphatidylcholine (LPC) into the cerebral WM of minipigs, which are not a distinct pig breed from the larger, landrace swine, but rather a variety of breeds that have been selectively bred to be smaller in size and which constitute some of the smallest known domestic swine (*Sus scrofa domesticus*). LPC microinjection provides temporal and spatial control, by inducing sequential de- and remyelination at predetermined locations(5, 7). Such control can minimize neurological deficits and, therefore, ethical constraints for CNS disease modelling – including in pigs, a species that is increasingly used for biomedical research, xenotransplantation or testing of novel human therapeutic antibodies(8–10). Neurological diseases that have already been modelled in the minipig subspecies include Parkinson’s disease, Huntington’s disease and experimental autoimmune encephalomyelitis (EAE)(11, 12),(13). Also, LPC and ethidium bromide-induced demyelination have been proven to lead to signs compatible with demyelination in the cerebral WM of larger, domestic landrace pigs(14) and to reliable demyelination of the rabbit cerebral WM(15) and of the non-human primate (macaque) optic nerve(16). However, there is considerable heterogeneity in the reported timelines of myelin-related events across species, ranging from initial reports of relatively quick demyelination (i.e., ensuing within 30 min and reaching full extent within 4 days(17)) and complete remyelination without evidence of substantial axonal damage within 5-6 weeks in the case of rodent models(18–20), to persistent demyelination conducive of axonal degeneration even 8 weeks after LPC injection in the macaque optic nerve. Interspecies differences such as the degree of collateral inflammation, the efficiency of lesion repopulation by oligodendrocyte progenitor cells (OPC), lesion volume, and reduction in oligodendroglial support to axons have all been postulated to contribute to the wide range of myelination and neurodegeneration responses(16).

For systematic and reliable placement of demyelinating lesions in the CNS of minipigs, we made use of computer-assisted navigation (CAN) as well as convection-enhanced delivery (CED), allowing for sufficient LPC penetration regardless of injection depth inside the tissue. CAN-technologies that rely on electromagnetic tracking systems(21) (EMTS) avoid imprecision due to instrument bending during surgery, since they derive the position and orientation of the instrument by measuring currents induced by an electromagnetic field using sensors at the instrument’s tip, rather than by extrapolation from imaging the shaft, as in optical tracking systems (OTS). The key advantage of the new EMTS we designed, compared to current techniques used in the human neurosurgery, is full compatibility with bendable instruments, because in addition to precisely measuring the tip position, we can also calculate the shape of the bend to detect and rectify undesirable deformations.

Using these neurosurgical innovations, we report here an EMTS platform for ***Mini****pig **S**tereotactic **W**hite-matter **I**njection using **N**avigation by **E**lectromagnetism* (*MiniSWINE*), which enables precise spatiotemporally controlled induction of demyelinating lesions in the minipig brain. We demonstrate longitudinal (i.e., at least 40 days post-induction) multimodal follow-up using diagnostic MRI protocols from human neuroradiology, ^11^C-PIB-PET, and neuropathological characterization as well as cross-validation against human inflammatory demyelinating disorders by MRI and electron microscopy (EM).

## Methods

A total of 9 young adult Aachen minipigs(22) (aged 19 ± 1 months at entry in experiment, mean ± s.e.m., life expectancy of 15-20 years), weighing 60.2 ± 2.32 kg (3 males and 6 females, mean ± s.e.m.) were used in this study. The study was performed in compliance with the EU Directive 2010/63/EU for animal experiments and the German Animal Welfare Act. All procedures were approved by the Ethics Committee for Animal Experiments of the Government of Upper Bavaria, Munich, Germany (Reference number ROB 55.2-2532.Vet_02-18-82). The animals were fed rationed feed rich in crude fibre. Water was provided ad libitum. We housed animals together, at least in pairs. Prior to the start of the study and daily after the start of the experiment, each animal was given a clinical examination by a veterinarian. The minipigs were acclimatised to the study procedures for at least two weeks.

### Experimental timeline

We performed an initial CT and Gd-enhanced MRI scan in general anaesthesia to aid surgery and injection trajectory planning, 10 ± 3 days before the first operative intervention. These were followed by 3 pairs of stereotactic injections in the cerebral WM, each pair placed in bilateral symmetric localization and performed 10 ± 3 days after the previous one. Immediately after the 2^nd^ pair of injections, as well as 10 ± 3 days after the 3^rd^ injection, we performed PET-MRI scans (Fig. 1a). The MRI scans, primarily, allowed for longitudinal follow-up of lesions over acute (10 days post-induction, dpi), intermediate (20 dpi) and subacute stages (40 dpi) and were performed according to standard sequence protocols recommended in the diagnosis of MS. The variability of the interscan intervals was mainly conditioned by scanner availability due to parallel use for humans in a clinical setting. The longitudinal PET scans were performed using ^11^C-PIB, to allow for assessment of myelination degrees of the lesions in acute and subacute stages. Euthanasia was performed up to 7 days after the last imaging procedure. Note that due to the use of radioactive tracer during the last imaging, dissection could not have been safely performed immediately after imaging, so that the animals were allowed time to recover after the general anaesthesia. Brain preparation and dissection followed directly after euthanasia.

**Figure 1.**
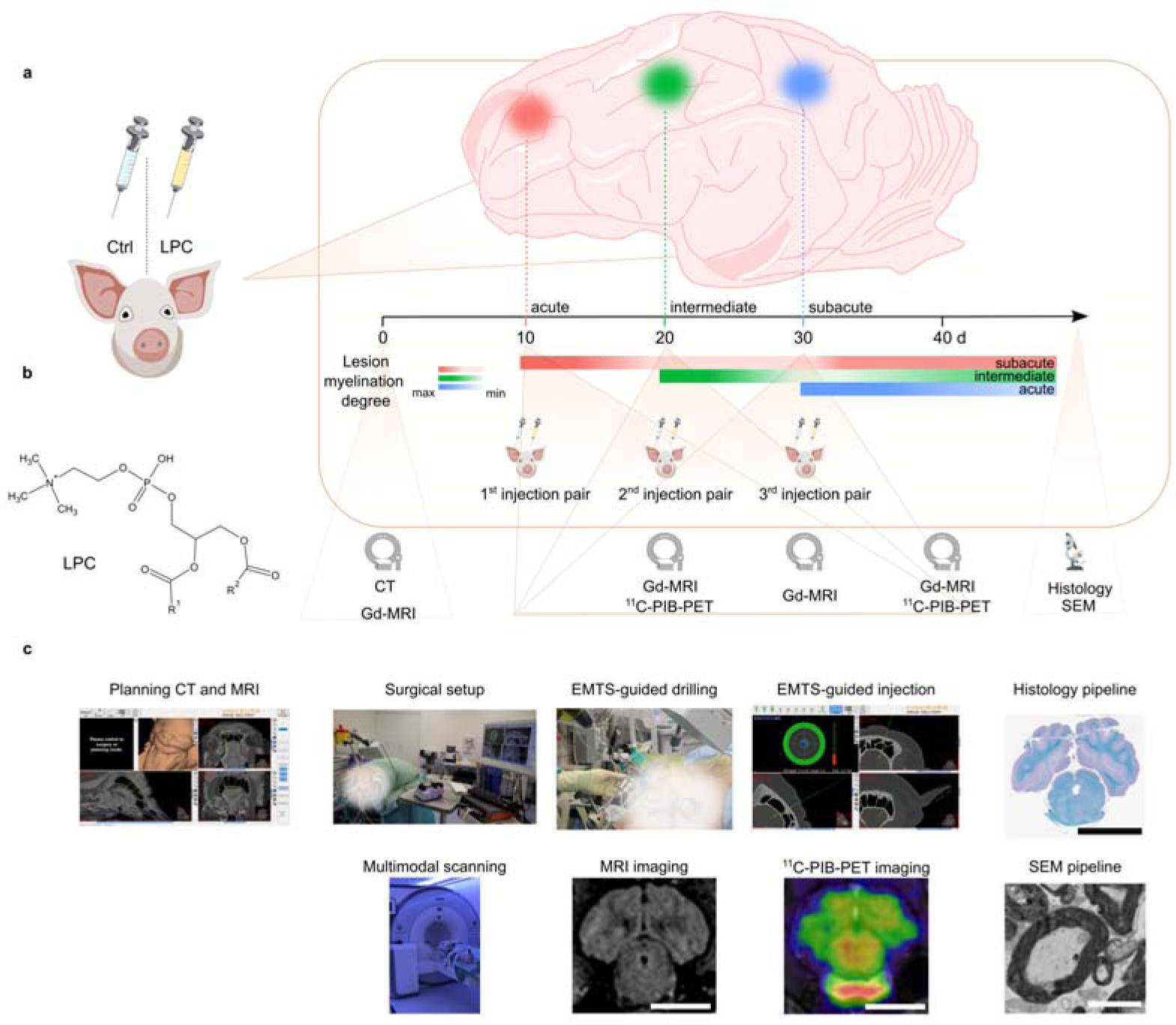
MiniSWINE methodology overview. a) Demyelination induction and remyelination follow-up timeline (d = days). Timepoint “0”: baseline imaging using CT and Gd-MRI. Timepoints “10-20-30 days post-injection”: every 10 days induction of a new demyelinating lesion at a distinct localization in the centrum semiovale and follow-up of the previous lesions. Gradient colour schematic reflects successive de- and remyelination stages. SEM = scanning electron microscopy. b) LPC chemical formula; R^1^, R^2^ = variable fatty acid chains. c) Overview photographs and photomicrographs of successive stages of *MiniSWINE*. White blurring obscures content with potentially strong emotional impact. Scale bar in insets “MRI imaging”, “^11^C-PIB-PET imaging” and “Histology pipeline” has 25mm, scale bar in inset “SEM pipeline” has 2.5µm. EMTS = electromagnetic tracking system.

### Preemptive imaging

CT scans were performed on a Siemens Symbia T6 scanner without contrast medium or intravenous contrast material and were essential for planning the surgical trajectory through the skull and the wide frontal sinuses of the minipigs. CT-image data were acquired in helical mode with a peak tube voltage of 130 kVp and a slice thickness of 1.25 mm. In order to avoid injuring macroscopical arterial and venous blood vessels when planning the injection trajectory, that would be only insufficiently represented in a CT scan, we added an intravenous contrast medium–enhanced MRI (Gd-MRI), undertaken using a field strength of 3T (Siemens Biograph mMR) and consisting of contiguous 0.5 mm T1 and 0.8 mm MPRAGE (Gd) sequences of a volume which included the skull and the whole snout of the animal (Fig. 1a, c).

### Lysolecithin preparation

For the intervention arm, lyophilized lysolecithin (Fig. 1b) in powder formulation (LPC, Sigma Aldrich) was dissolved in iso-osmolar phosphate-buffered saline (PBS) at 40°C achieving a concentration of 20mg/ml. The solution was aliquoted and stored at -20°C, then thawed just before use. For the control arm, we used only sterile PBS.

### Injection system

In order to achieve efficient (reflux-free) and precisely placed and dosed CED(23) of LPC in the intervention and PBS in the control arm, we designed a special electromagnetic tracking-system with a compatible cannula (Fig. 2a, b). The cannula had a rounded tip, was made of stainless steel and featured a 3D-printed biocompatible Luer-Lock fitting (Fig. 2c, d). It had an outer diameter of 2.0 mm, an inner diameter of 0.85 mm, a total length of 250 mm and was designed to accommodate 20 G epidural catheters (Braun Perifix, Germany). By inserting a sensor wire designed and manufactured by us into the cannula and connecting it to the Luer-Lock fitting, the cannula could be tracked and visualized via the navigation system (Fig. 2b, c). The sensor wire had a Luer-Lock, a diameter of 0.8 mm and was inserted about 1 mm short of the tip of the cannula. An electromagnetic sensor was integrated into the body and tip of the wire, allowing not only tracking of the position and orientation of the wire tip, but also measurement of the bending of a flexible instrument and detection of defects in rigid instruments.

**Figure 2.**
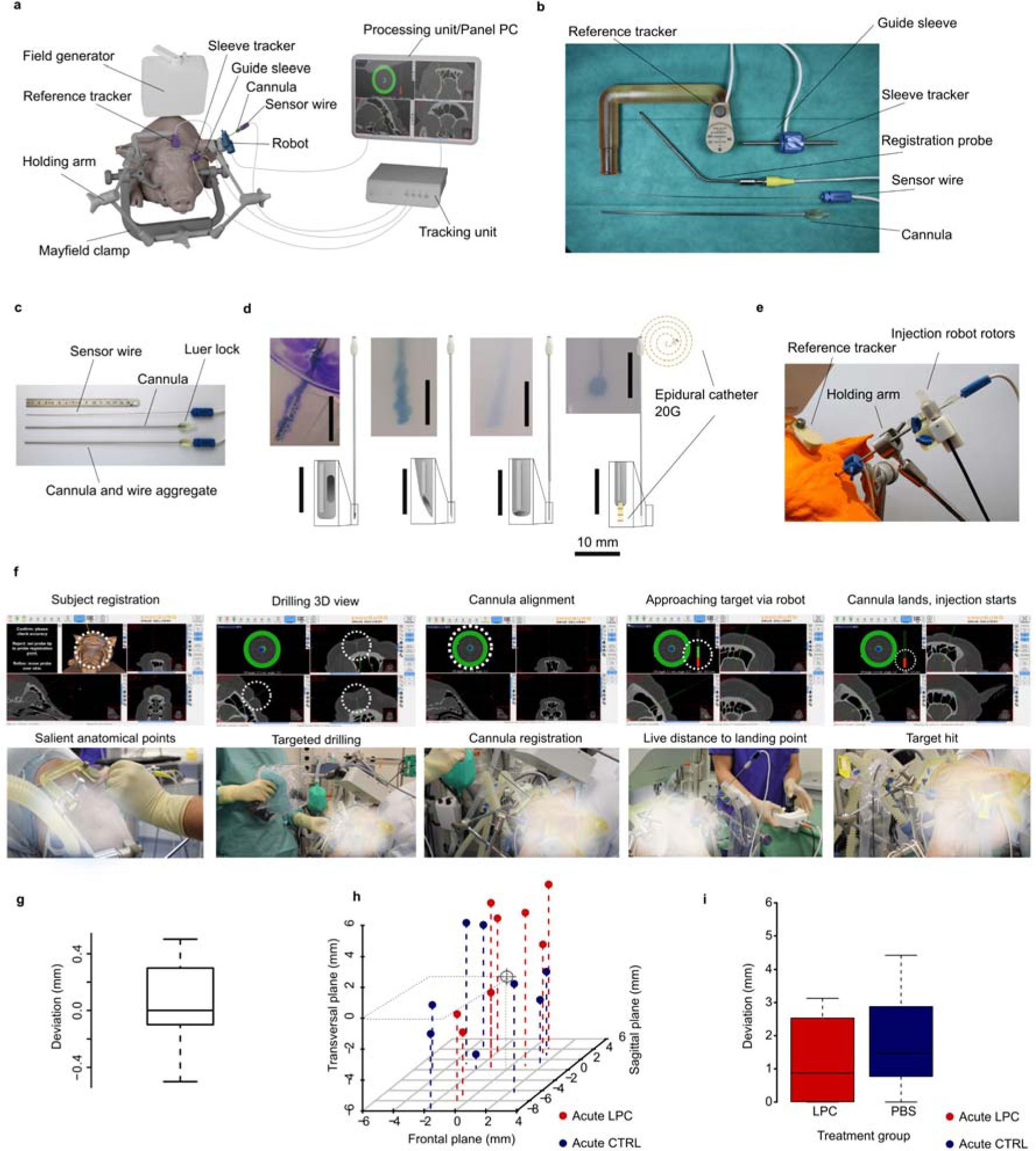
MiniSWINE: electromagnetic-tracking system (EMTS) and surgical procedure. a) *MiniSWINE* setup for EMTS b) The reference tracker was rigidly attached to the minipig forehead. The tracked guide sleeve was essential for precise alignment of the drill. c) Cannula assembly. The CED cannula was trackable as long as it contained the sensor wire. Once on target, the wire was replaced with the single use 20G epidural catheter. d) Sequential cannula development, cannula tip profiles, and empirical spread functions of CED in 0.2%-0.6% agarose gel. Evolution from a modified human brain biopsy cannula with a side-opening near the tip (far left) up to our definitive “telescopic” design (far right) with a round bevelled edge. Notice the near-ideal spherical spread function of the latter. e) Injection “hands-on” robotic system assembly consisting of a control unit and a small robot for the controlled and tracked insertion of the cannula. f) *MiniSWINE* software in operator view, 1^st^ row and corresponding surgical overview, 2^nd^ row. White dotted circles in 1^st^ row mark system feedback to the operator. White blurring obscures content with potentially strong emotional impact. g) Mean *in vitro* deviations from target in a phantom test, of 0.23 ± 0.03 mm (mean ± s.e.m., n = 25 trials). h) Scatterplot of 3D deviations from target *in vivo* in all 3D planes; black “target” set in the common origin of the coordinate system. i) Mean *in vivo* deviations from target by study group: no significant difference between acute LPC lesions (1.24 ± 0.5 mm, mean ± s.e.m., n = 8) and acute CTRL lesions (1.83 ± 0.5 mm, mean ± s.e.m., n = 8), 1-way ANOVA with *df* = 1, *F* = 0.71, *p* = 0.41.

The injection cannula along with the sensor wire were inserted with the aid of the robotic system mounted on the guide sleeve and controlled via a joystick by the operator. The robot was connected to the navigation system, from which it received positional data. This data was used to stop the controlled insertion of the injection cannula at the instance the desired injection depth was reached. For minimising mechanical damage to the tissue at the injection site due to insertion of the cannula, we employed a 20 G epidural catheter that protruded around 3-5 mm from the tip of the cannula. In this way, we achieved a telescopic profile at the tissue-cannula interface, which has been shown to minimise reagent reflux(24) (Fig. 2d, e; Suppl. Video 1). Per instance, 250 µl of LPC (intervention hemisphere) or PBS (control hemisphere) were injected over 50 min (5 µl/min) using a Harvard microinjector (Harvard Pump 11 Elite, Harvard Apparatus, USA), the slow rate being necessary to allow for diffusion in the cerebral parenchyma and to avoid significant backflow, i.e., reflux of reagent(23), the injection cannula remaining in place for 10 min after complete reagent delivery, before it was extracted (total tissue insertion time 60 min) (Fig. 2f).

### Navigation system

The navigation system, designed and fabricated by Ergosurg GmbH, consisted of a processing unit with a navigation software which was connected to an electromagnetic tracking unit (Fig. 2a). This electromagnetic tracking unit generated a defined alternating electromagnetic field imparting a discretised measuring volume to the surgical area. In this electromagnetic field, the position and orientation of instruments with integrated sensors connected to the tracking unit could be measured. As such, after registration of the salient surface features of the minipig head, instruments could be visualized relative to the subject in pre-surgical CT and/or MRI image data in real time. As a reference point for this registration a so-called reference tracker had to be rigidly attached to the head (Fig. 2b). For the registration itself a navigated probe for scanning the skin and probing landmarks was connected to the system. For precise alignment of the guide sleeve to the trajectory and its entry point under navigation, we built a sensor clip and attached it to the sleeve. The sensor wire was also connected to the tracking unit.

The navigation software we developed (Drug Delivery V6.3.1) supported importing and the fusion of CT and MRI data, planning of trajectories and entry points, surface registration and real-time visualization of instruments/cannulas in canonical planes (axial, coronal, sagittal) as well as in 3D segmentations. In addition, the software supported the control of a small medical robot system we designed for navigated insertion control of the cannula. During surgery, the navigation software displayed real-time feedback on the position and orientation of the probe, of the trackable inner cannula, and the deviation from the planned trajectory and the injection target, while concomitantly projecting the imaging data and the virtual 3D model on a screen, as long as the sensor wire was contained in the lumen of the cannula. In our paradigm, salient anatomical maxillofacial land points (i.e., nasal ridge, frontal bone, zygomatic arcades, mandibular joints) were used as landmarks to guide the registration probe. These allowed for pre-emptive evaluation of accuracy with the trackable probe. The field generator was positioned in a consistent position between trials near the animal’s head so that the head and, most importantly for the stereotactic coordinates, the vertex were in the centre of the electromagnetic measuring volume.

### Surgical procedure

Prior to anaesthesia, the pigs were examined by a veterinarian. All animals were clinically healthy. The basic anaesthesia regime for both surgeries included sedation by combination of ketamine 15 mg/kgbw, azaperone 2 mg/kgbw and atropine 0.1 mg/kgbw intramuscularly. An intravenous catheter was placed in the lateral auricular vein. Induction of anaesthesia by propofol (ca. 4-8 mg/kgbw titrated to effect) intravenously (IV) to achieve an adequate depth of anaesthesia for oral intubation with a 5-7 mm cuffed endotracheal tube. Anaesthesia was maintained by a continuous IV infusion of propofol 2% (ca. 2.5-7 mg/kgbw/h). Minipigs were mechanically ventilated in volume-controlled mode and oxygenated with 1-2 L O2 / min, mixed with 1 L atmospheric air / min. Tidal volumes (ca. 10 ml/kg) and breathing rate (ca. 12/min) were adapted to the end-tidal CO2. PEEP was kept at 5mmHg. Saline was infused IV at a maintenance rate of ∼10 ml/kg/hour. Intraoperative monitoring included reflex status, heart rate and peripheral arterial oxygen saturation (SpO2) by use of a pulse oximeter, electrocardiogram, end-tidal CO2 and rectal body temperature. External warm-air supply was provided. A single-shot Cefuroxim IV (500-750 mg) was administered before skin incision. For multimodal analgesia during the surgery, metamizole (50 mg/kgbw), carprofen (4 mg/kgbw) and fentanyl-boluses (0.001 – 0.01 mg/kg) every 20-30 minutes were administered IV. For the imaging procedure, a second peripheral venous catheter was placed into the V. antebrachia or V. cephalica for administration of the radioactive tracer.

Minipigs were positioned prone, followed by shaving and disinfection of the operative situs. The head was fixed in a Mayfield clamp (Pro Med Instruments, Freiburg, Germany) (Fig. 2a, f). Using a probe, the surface of the skull was registered and thus virtually superimposed on the planning imaging dataset (a fusion between CT and MRI) (Suppl. Video 2). A trackable guide sleeve was aligned to the planned trajectory during real-time feedback from the navigation software. Once the intended trajectory was achieved, the trackable guide sleeve was fixed in place and a 2.6 mm-wide burr hole trepanation was drilled through the lamina externa and interna of the frontal bone, until the dura mater (haptic loss of resistance) was reached (Fig. 2f, Suppl. Video 3).

Then a medical robot was attached at the entry of the guide sleeve and the injection cannula along with the inner trackable wire were inserted manually into the robot and into the guide sleeve. With the feedback of the navigation system the robot then inserted the cannula into the brain and stopped automatically when the intended intraparenchymal target was reached (Suppl. Video 2). Finally, the sensor wire was removed and replaced with the 20 G epidural catheter, through which, after connection to the Harvard microinjection pump, 250 µl LPC or 250 µl PBS at a rate of 5 µl/min were infused over 50min.

We aimed at positioning roughly equidistant LPC/PBS injections at least 5mm apart in rostro-caudal succession in the centrum semiovale of both hemispheres. Post-hoc quantification of planned landing targets gave following mean injection coordinates computed with the vertex as a reference: -2.8 ± 0.7 mm (mean ± s.e.m.) and 13.2 ± 2.2 mm (mean ± s.e.m.) in the axial plane, -3.6 ± 1.7 mm (mean ± s.e.m.) in the coronal plane and in a rostro-caudal gradient in the sagittal plane, beginning with the first pair of injections at -53.1 ± 1.8 mm, the second pair of injections at -41.9 ± 0.6 mm (mean ± s.e.m.), and the third pair of injections at 35.62 ± 1.1 mm (mean ± s.e.m.). The variability of the injection coordinates is conditioned, in our view, by two main factors. Firstly, in contrast to rodent brain LPC injections, where invariable stereotactic coordinates relative to an anatomical structure (e.g., bregma) can be formulated and translated between individuals(25), in minipigs, due to the larger brain volume (approximately 70-80 000 mm³ in our cohort), there is the necessity of avoiding vasculature of more than 0.8mm diameter in the injection trajectory, which naturally differs up to a certain extent in anatomical distribution among individual minipigs. Secondly, there is inherent variability in the minipig brain size, which, even if it were equivalent to the extent of variability between adult mice (volume of 400-500 mm³ according to the literature(25)), due to the difference in scale makes even subtle discrepancies relevant and requires adaptation of injection targets for each individual minipig.

After injection, the incision was sutured. Total surgical time amounted to approximately 3 h, from which 2 h were represented by the total injection time, while around 30 min were required for mounting the head in the Mayfield clamp and for navigation.

Postoperative analgesia was administered in the form of daily carprofen PO or IM (4 mg/kgbw) over 5 days as well as buprenorphine IM (0.01-0.1 mg/kgbw) for 3 days. Antibiosis was maintained in parallel with daily amoxicillin PO or IM (15 mg/kgbw). Veterinarian clinical observations were performed once daily for the entire duration of the study, in order to ascertain that the animals did not develop any neurological deficits as a consequence of the intracerebral injections.

### Longitudinal MRI imaging

Longitudinal MRI imaging consisted of scans performed immediately after the 2^nd^ and 3^rd^ pairs of stereotactic intracerebral injections, as well as 10 ± 3 days after the 3^rd^ pair of injections. Minipigs were placed prone in a 3.0 T Siemens Biograph mMR (Siemens, Germany) with the head rigidly fixed in a standard cage head coil.

Concordant to the 2021 MAGNIMS consortium(26) recommendations on the use of MRI in patients with MS, we performed sagittal 3D T2-weighted (1.5 mm slice thickness, echo time = 107 ms, repetition time = 4220 ms), sagittal and axial isotropic 3D T2FLAIR (1 mm slice thickness, echo time = 393 ms, repetition time = 6000 ms, inversion time = 1800 ms), T1-weighted post-contrast/T1Gd (contrast medium: Gd, Dotagraf, 0.5 mmol/ml, 2 mm slice thickness, echo time = 2.84 ms, repetition time = 250 ms), T1-weighted MP RAGE(27) (1 mm slice thickness, echo time = 3.13 ms, repetition time = 1920 ms, inversion time = 999 ms), double-inversion recovery for detecting potential juxtacortical lesion extensions (DIR, 1 mm slice thickness, echo time = 318 ms, inversion time = 3000 ms, repetition time = 7500 ms), T1-weighted “black-blood” (slice thickness = 0.5 mm, echo time = 11 ms, repetition time = 1070 ms) and, in addition to the MAGNIMS sequences, susceptibility-weighted imaging (SWI, slice thickness = 9.6 mm, echo time = 20 ms, repetition time = 28 ms).

### MRI image analysis

The images were processed with the software MITK Workbench v2018.04.2 (www.mitk.org) using its 3D segmentation toolbox. Regions of interest (ROIs) were semi-automatically segmented in T2FLAIR, T1Gd and SWI sequences, and their volumes (in mm³), excluding the injection trajectory path, were quantified. Volumes from the LPC injection site were compared to their contralateral control counterparts of the same stage (PBS injection). Deviations from the intended placement target *in vivo* were determined via automatic superposition of CT and MRI datasets from the baseline imaging with the ones performed after the corresponding lesion placement, from which the mean superposition imprecision between the CT and MRI dataset was subtracted. The superposition imprecision was determined at the midpoint of the sphenoid bone ridge, which was reliably identifiable both in CT and T1 MRI.

As in the case of biopsies in the context of neurosurgical patients, every insertion of a cannula in the cerebral parenchyma, regardless of diameter, induces SWI susceptibility corresponding to (asymptomatic) microhaemorrhage along the injection trajectory(28). We excluded from the analysis lesions which exhibited a volume of SWI hyperintensity higher than 100mm³, a threshold derived by considering the volume (*V*) of a cylinder of microhaemorrhage given by the outer diameter (2mm, see Methods, 2 × *r*) and length (250mm, *h*) of the cannula (*V* = π *x r*^2^ *x h*). In this way, we excluded microhaemorrhages of dimensions beyond the lumen of the cannula: thus, 20 out of initially 80 lesions (25%) were excluded from the analysis (Suppl. Fig. 1).

### Longitudinal PET imaging

PET studies were performed immediately after the MRI after the 2^nd^ pair of stereotactic intracerebral injections, as well as 10 ± 3 days after the 3^rd^ pair of injections. For PET scans we used ^11^C-PIB as a tracer, originally developed for amyloid plaque tracking, but pioneered in the context of myelin tracing in a proof-of-concept study on 2 MS patients(29), produced according to standard protocols(30, 31). After the surgery, outside the scanner, animals were injected through an intravenous catheter placed in the lateral auricular vein with around 370 MBq of ^11^C-PIB. Between 45- and 60-min post-injection, a 15-min list-mode scan was acquired in 3D mode on the Siemens Biograph mMR as well as decay of radioactivity, and smoothed using a Gaussian kernel.

Emission sinograms were reconstructed using ordered subset expectation maximization (OSEM) corrected for attenuation, scatter, randoms, dead time as well as decay of radioactivity, and smoothed using a Gaussian kernel. Mean standardized uptake values (SUV) were quantified and compared using the software Weasis v.3.8.1 (https://nroduit.github.io/en/) after manual delineation of lesion volumes of the same stage slice-by-slice on the intervention and control sides in the overlapped PET/MRI datasets and calculation of SUV values by area-weighted averaging.

### Histopathology

With a latency of 1-7 days following the 3^rd^ PET/MRI scan, animals were euthanized by pentobarbital overdose during deep anaesthesia, a total of 49 ± 7 days from the initial pair of intracerebral injections. The latency between the 3^rd^ PET/MRI scan and euthanasia was conditioned by persisting radiotracer-related radioactivity precluding brain dissection immediately after the PET scan. After dissection, the brains were fixed in 4% PFA via submersion. We embedded brains in paraffin wax which was melted in a constant temperature oven at 64°C and kept at 60°C after complete melting, according to previously published procedures(32). The cooled (4°C) paraffin blocks were then sliced in coronal orientation at a thickness of 4-5 µm on a semiautomated rotary microtome (Leica, Germany). These sections were then collected, mounted and stained by Luxol fast blue (LFB, Klüver-Barrera) – periodic acid-Schiff (PAS), as well as haematoxylin and eosin (H&E).

For immunohistochemistry, 2 μm thick slides were cut with a standard microtome and dried at 76 °C for 30 min. Glial fibrillary acidic protein (GFAP), ionized calcium-binding adaptor molecule 1 (Iba1), oligodendrocyte transcription factor 2 (Olig2), neuronal nuclear antigen (NeuN), cluster of differentiation 3 (CD3) and CD68 immunostaining was performed using a fully automated staining system (BOND-III; Leica Biosystems, UK). In brief, the slides were pretreated in pH 8.4 buffer for heat-induced epitope uncovering and H_2_O_2_ as inhibitor of endogenous peroxidase in order to prevent unspecific bindings of the primary antibody. Afterwards, the slides were charged with anti-GFAP (monoclonal, rabbit, dilution 1:500; Cell Marque, Rocklin, CALIF, USA), anti-Iba1 (polyclonal, mouse, dilution 1:1000; Fujifilm Wako, Japan), anti-Olig2 (monoclonal, mouse, ready to use; Cell Marque, Rocklin, CALIF, USA), anti-NeuN (monoclonal, mouse, dilution 1:1000; Cell Marque, Rocklin, CALIF, USA), anti-CD3 (monoclonal, rabbit, ready to use; Roche, Switzerland), or anti-CD68 (monoclonal, mouse, dilution 1:200; Cell Marque, Rocklin, CALIF, USA) antibodies. For antibody detection and counterstaining, the BOND Polymer Refine Detection Kit (Leica Biosystems, UK) was used. Positive controls were used as quality assurance.

Images were processed using Fiji ImageJ (https://imagej.net/software/fiji/)(33). Olig2+, NeuN+ and CD3+ cell numbers were manually quantified per 300 × 300 µm²-sized region of interest (ROI), while activated astrocytes and reactive microglia were manually quantified as a proportion of a 1 × 1 mm² ROI. We refrained from CD68 quantification because of low signal-to-noise ratio, presumably due to low cross-species primary antibody affinity. The stained sections were then mounted and examined under a bright field microscope (Leica, Germany), then scanned and uploaded using a microscope-mounted camera and the software PathoZoom Slide Cloud (https://www.pathozoom.com/). Demyelination areas were quantified in LFB-stained slides and compared between intervention (LPC) and control (PBS) study groups from lesions of the same stage using Slide Cloud and Fiji ImageJ (https://imagej.net/software/fiji/)(33). Since lesions were relatively small and regularly shaped compared to the brain volume across all three dimensions, we calculated demyelination areas by manual tracing in Image J and summation of areas in analogy to the rectangular method of estimation of morphometric volume(34, 35).

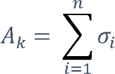

where *A_k_* represents the area of lesion *k, n –* the number of sections obtained for lesion *k,* and σ – the cross-section area in mm² of section *i*. We opted to not compute volumes, since this would, due to the simple multiplication between the slice thickness and section-by-section area, provide no additional information (derivative of a constant is null), but risk introducing additional measurement errors inherent to deviations in slice thickness or regularity of slice sampling during cutting.

For immunofluorescence, the slices were deparaffinized and rehydrated by emerging the slices in xylene and series solutions of decreasing concentrations of ethanol. The antigen retrieval was carried out with the All-Antigen Unmasking Kit from Bio Legend following the manufacturer’s protocol. The sections were then blocked with 5% goat serum for 2 hours at room temperature. Afterwards, they were incubated overnight with primary antibodies, followed by secondary antibodies. After incubation with secondary antibodies, the sections were mounted with Prolong Gold Antifade mounting media with DAPI from Invitrogen. We used antibodies against GFAP (glial fibrillary acidic protein, Cat#13-0300, ThermoFisher), MOG (myelin oligodendrocyte glycoprotein, Cat#MAB5622, Merck Millipore) and NFH (neurofilament heavy chain, Cat#AB5539, Merck Millipore). We used Alexa Fluor 555nm labelled goat anti-mouse secondary antibody (Cat#A28180, ThermoFisher) against MOG, Alexa Fluor 488nm labelled goat anti-rat secondary antibody (Cat#A11006, ThermoFisher) against GFAP and Alexa fluor 488nm labelled goat anti-chicken secondary antibody (Cat#A11039, ThermoFisher) against NFH. The stained sections were then mounted and examined under a confocal microscope (Olympus FV1000) and scanned using Olympus Fluoview Software. We refrained from the quantification of immunofluorescence intensities or immunofluorescent areas and resorted to a descriptive analysis because we observed a degree of crosstalk between the GFAP- and MOG-antibodies.

### Scanning electron microscopy

Minipig brain as well as human biopsy tissue used for the scanning electron microscopy (SEM) experiment was immersed directly after dissection in fixative solution (2.5% glutaraldehyde, 4% paraformaldehyde in phosphate buffer, 2mM calcium chloride, 0.1 mM cacodylate buffer). After fixation for 48 h at 4°C, the lesion areas (intervention or control, same lesion stage) were washed in 0.1 M cacodylate buffer and dissected. We applied a standard rOTO protocol *en bloc* staining(36) including postfixation in 2% osmium tetroxide (Science Services), 1.5% potassium ferricyanide (Science Services) in 0.1 M sodium cacodylate (Science Services) buffer (pH 7.4). Staining was enhanced by reaction with 1% thiocarbohydrazide (Sigma) for 45 min at 40°C. The tissue was washed in water and incubated in 2% aqueous osmium tetroxide, washed and further contrasted by overnight incubation in 1% aqueous uranyl acetate at 4°C and 2 h at 50°C. Samples were dehydrated in an ascending ethanol series and infiltrated with LX112 (LADD). SEM was performed on 100 nm thick sections collected onto carbon nanotube tape (Science Services) and mounted onto a silicon wafer (Microchemicals). EM micrographs were acquired on a Crossbeam Gemini 340 SEM (Zeiss) with a four-quadrant backscatter detector at 8 kV. In ATLAS5 Array Tomography (Fibics), wafer overview images (1.5 µm/pixel) as well as medium (200 nm) and high (10-20 nm) resolution images were acquired. Images were processed using the software Fiji(33) (https://imagej.net/software/fiji/), while manual assessment of myelination degree in SEM images used the well-established g-ratio(37). Quantification was performed on 10-20 nm resolution images for presence/absence of (myelinated) axons per area. We counted a total of 1499 axons in the minipig subcortical WM across all stages and 1614 axons in the human samples.

### Ethics and human samples

Written informed consent was obtained from each participant treated in the Department of Neurology, Klinikum rechts der Isar, Technical University of Munich, Germany. Samples were collected as part of the biobank project of the Department, which pertains to the Joint Biobank Munich in the framework of the German Biobank Nodes. Patient A (*acute, aHum*), a 19-year-old female needed a diagnostic stereotactic biopsy from the clinically manifest cerebellar white matter lesion and was later diagnosed with elapsing-remitting MS. Patient S (*subacute, sHum*), a 68-year-old male, had a diagnostic stereotactic biopsy from the right parietal subcortical white matter ascertaining the diagnosis of ADEM at a subacute stage. This was taken after an initial positive response to *ex juvantibus* immunosuppressive therapy (1 g methylprednisolone intravenously per day over 5 days), therefore providing time for incipient remyelination of the biopsied plaque.

### Statistics

A minimal number of minipigs and induced lesions was predetermined by statistical power analysis. In order to achieve an intraindividual correlation of *r*²= 0.85, we needed at least 3 animals with at least 3 × 3 = 9 samples to a two-sided statistical significance level of 5% (α = 0,05), considering a previously described attrition rate of 50% in the literature(38). Due to the nature of longitudinal lesion follow-up and consecutive lesion induction in the same animal, our sampling power has been highest for the acute stage (n = 22 lesions in 8 animals) and lowest in the subacute stage (n = 6 lesions in 5 animals). The spatial and temporal separation of lesions allowed them to be regarded as independent statistical units. Statistical analysis was performed using the open-source software R (https://www.R-project.org/, version 4.1.0) and R Studio (version 1.4.1717). Data were tested for normality and equality of variance using the Shapiro-Wilk test and, respectively, the Bartlett test. If the data passed normality and homoscedasticity criteria, we performed one-way repeated-measures ANOVA tests with Bonferroni adjustments for multiple comparisons. If they did not, then we performed the Kruskal-Wallis test with Dunn’s multiple comparisons test. All statistical tests were two-sided and performed to a significance level of p <= 0.05.

## Results

For the *MiniSWINE* methodology (Fig. 1a), we resorted to stereotactic microinjection of LPC (Fig. 1b) for optimal spatio-temporal control. We chose EMTS (Fig. 1c, Fig. 2a) over OTS, as EMTS does not require a visual axis (i.e., line-of-sight). Moreover, as EMTS does not rely on direct visualization, the handling of the navigated drill guide on small surfaces such as the porcine forehead is easier. Finally, with EMTS navigation, the porcine head positioning in the clamp holder does not have to be adjusted intraoperatively, which can be the case when faced with line-of-sight issues in OTS.

### Stereotactic navigation system via electromagnetic tracking

The navigation system that we designed and fabricated consisted of a processing unit/panel PC running our navigation software, which was connected to an electromagnetic tracking unit, ETU (Fig. 2a). By means of a planar field generator, the ETU created a defined alternating electromagnetic field imparting a measuring volume of approximately 0.5 m × 0.5 m × 0.5 m to the surgical area(39). The field generator was mounted on a holding arm and positioned near the animal’s head, so that the surgical area was in the centre of the EMTS’s measuring volume. In this electromagnetic field, the position and orientation of instruments with integrated sensors connected to the ETU could be measured once a registration process was completed. The registration required a reference point with a fixed position relative to the minipig’s head, for which we firmly attached a reference tracker to the head via the holding arm (Fig. 2b). The registration itself was based on surface matching, where the system measured the tip of a navigation probe with which the operator touched salient surface features of the minipig head, that were then matched to the image data by our software (Suppl. Video 2). This surface matching took 15 ± 0.1 s (mean ± s.e.m.) *in vitro* (for 110 points) and 96 ± 4.4 s (mean ± s.e.m.) *in vivo* (for 482 points), and thus did not impact on surgical time significantly.

To guide the drill and the injection cannula, we attached a guide sleeve to another holding arm, which carried a sleeve tracker for precise alignment. The guide sleeve with its 6 degrees of freedom (DoF) tracker (Fig. 2b) was positioned by the holding arm which was attached to the Mayfield clamp supporting the head of the minipig. For the injection itself, a custom-made cannula with a Luer-Lock fitting was fabricated. For navigating the cannula, we fabricated a so-called sensor-wire, a superelastic nickel titanium alloy wire with a 5 DoF sensor in the tip, which could be inserted into the cannula and attached via the Luer-Lock fitting (Fig. 2c).

We determined empirically which cannula tip profile resulted in the optimal spread patterns of the injected substance by *in vitro* injections into 0.2% or 0.6% agarose gels (which have been described to imitate the consistency of the mammalian brain(24, 40)). We tested volumes and infusion rates compatible with CED(41), i.e., 250µl at 5 µl/min. Known problems of the standard needle profiles used in stereotactic biopsy needles, which have a bevel or side aperture near the tip, are backflow and an elongated spread pattern along the needle tip. It has been shown previously that backflow can be influenced by the consistency and the profile of the needle tip(42). Therefore, we custom-designed an autoclavable cannula of 2 mm outer diameter with a rounded tip (Fig. 2c), which minimised haemorrhage risk. Through this needle a standard single-use 20G silicon epidural catheter (0.85 mm outer diameter) could be inserted, replacing the sensor wire, so that it protruded 3-5 mm from the tip. In this way, the needle tip was not only soft, but exhibited a step-profile which has been described to minimise backflow issues(42). Consequently, the spread functions reached elliptical/almost spherical profiles of at least 5 mm radius, which was the minimal spatial resolution of the PET scanner (Fig. 2d).

We developed a “hands-on” robotic system consisting of a control unit and a small robot for the navigated insertion of the cannula (Fig. 2e). The control unit was connected to the navigation system and could be mounted on any operating table. For control of the robot by the navigation or by hand, the control unit had a joystick and a safety button preventing unintentional activation. The robot was in turn connected to the control unit and could be mounted on the guided sleeve. It had two steel shafts with sterilisable silicone covers for inserting the navigated cannula via the guided sleeve into the brain. Only one of these shafts was motor driven, so that manual insertion or removal of the cannula was possible at any time.

Care was also taken to optimize the flow of the EMTS-assisted surgical procedure, by providing the essential visual feedback quickly via our software, in correspondence with the haptics sensed by the surgeon. After registration of the minipig head, we could visualize trackable instruments relative to the brain in pre-surgical CT and/or MRI image data in real time in the 3 orthogonal planes (axial, coronal, sagittal) and in 3D segmentations. In addition, the software showed the remaining distance of the cannula’s tip to the target. The planning of the trajectory could be done in the planning tool integrated in the navigation software or separately on a laptop or PC with the navigation software installed. For confirming registration accuracy, the surgeon touched anatomical landmarks of the head with the tip of the probe and checked the visualized position, revealing a registration error of 0.55 ± 0.01 mm *in vitro* and 0.87 ± 0.013 mm *in vivo* (mean ± s.e.m.). For controlling the movement and insertion depth of the navigated cannula, the cannula was inserted into the guide sleeve and thus into the brain by the “hands-on” robotic system or, alternatively, manually by the surgeon. For the robotic insertion, the robot was also mounted to the sleeve and could move the cannula via its motor-driven shaft with a speed of 11 mm/s and a movement resolution of <0.1 mm. The surgeon could control the robotic system via the navigation software, which provided visual feedback of the remaining distance to target on a screen for the operator and stopped the robot / the cannula’s movement automatically as soon as the tip of the cannula reached the target (Fig. 2f, Supplementary Videos 1,3).

With a minipig phantom head, 3D printed from a CT dataset of one of our animals, in a typical operating room setup, an *in vitro* accuracy with a mean positional error of 0.23 ± 0.03 mm (mean ± s.e.m.) was achieved (Fig. 2g). We have already demonstrated *in vivo* that the system level accuracy of the navigation system exhibits a mean positional error of ≤ 2.0 mm and a mean trajectory error of ≤ 2° in a human spinal surgery setting(43). In our case, the *in vivo* injection imprecision in our *MiniSWINE* brain surgery setting did not discriminate between the 3 orthogonal planes (Fig. 2h), and averaged 1.54 ± 0.4 mm (mean ± s.e.m.) over both study groups, with no significant bias towards LPC or CTRL (Fig. 2i). This was sufficient considering the necessary spatial resolution of the detection systems employed and the anatomy of the minipig brains structures, which have a similar configuration to, but are even more compact than human ones.

### MR-imaging of de- and remyelination

A reliable large animal model for an inflammatory CNS disorder such as MS would need to be amenable to diagnostic imaging on a scale similar to human patients. Standardised MRI scan protocols should include axial T2-weighted, sagittal and axial T2-weighted (preferably 3D) FLAIR, post-contrast axial (or 3D sagittal) T1-weighted, high-resolution T1-weighted (isotropic 3D acquisition), double inversion recovery (DIR) for the diagnosis of spatial and temporal dissemination of demyelinating (brain) lesions in MS, as defined by the 2021 MAGNIMS consortium(26) and the 2017 McDonald criteria(44) for the diagnosis of MS. One diagnostic method with potential for the specific quantification of de- and remyelination *in vivo* and therefore evaluation of future remyelinating treatments is PET employing WM-binding tracers, such as the thioflavin-T derivative 2-(40-methylaminophenyl)-6-hydroxybenzothiazole (Pittsburgh compound B, PIB), pioneered in a proof-of-concept study on 2 MS patients(29). To the MRI scan protocol derived from the MAGNIMS criteria, we added SWI (susceptibility-weighted imaging) sequences, that are particularly sensitive to iron content from venous blood and haemorrhage, in order to ascertain that tissue changes after LPC/CTRL-injections are not due to injection-induced haemorrhage.

First, we investigated whether the MRI signatures of the LPC-induced lesions matched expected patterns on established protocols developed and used in clinical practice for MS patients. We performed MRI imaging at intervals designed to capture critical stages in lesion evolution, as known from LPC-induced demyelination in rodent models(17) (Fig. 3a). We investigated an acute stage at a median of 9 days (IQR 7-14 days, *n_aLPC_* = 22 *aLPC* lesions and *n_aCTRL_* = 22 *aCTRL* lesions), an intermediate stage at a median of 20 days (IQR 19-23 days, *n_iLPC_* = 12 *iLPC* and *n_iCTRL_* = 12 *iCTRL* lesions) as well as a subacute stage at a median of 30 days (IQR 28-33 days, *n_sLPC_* = 6 *sLPC* and *n_sCTRL_* = 6 *sCTRL* lesions) after lesion induction. Note that by longitudinal imaging in the same animal, lesions at earlier stages accumulated in higher numbers than lesions at latter stages, so that we obtained a 3 : 2 : 1 ratio between the number of acute : intermediate : subacute lesions in our study (Suppl. Fig. 1), that we could then examine multimodally.

**Figure 3.**
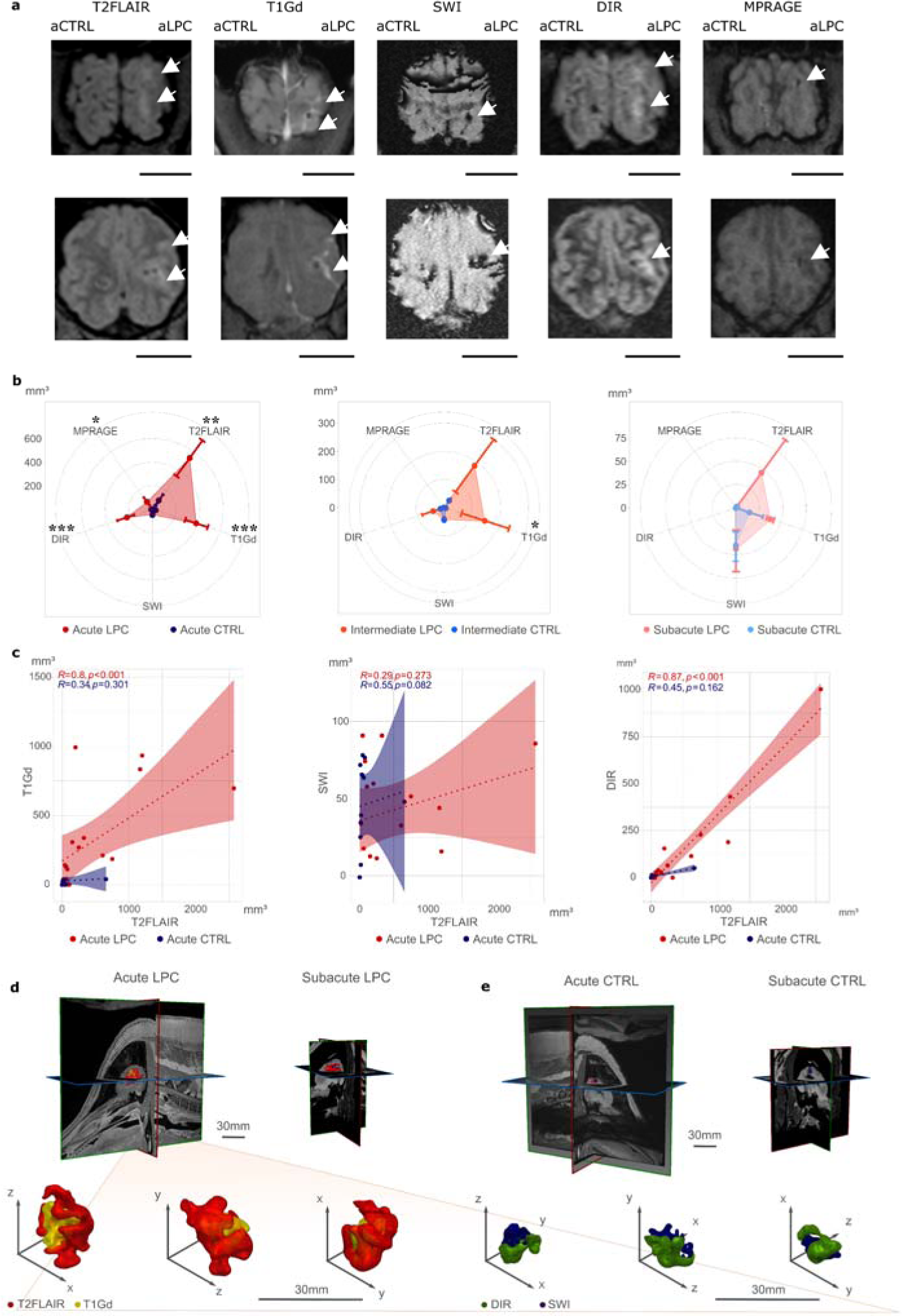
MiniSWINE MRI. a) Example axial tomographic planes, each row corresponding to one acute lesion from a separate minipig, scale bar = 2 cm. b) Mean ± s.e.m. MRI signals in each stage stage (*aLPC* n = 18, *aCTRL* = 13; *iLPC* n = 11, *iCTRL* n = 9, *sLPC* n = 4, *sCTRL* n = 5). Statistical significance and *p* values were determined by the Kruskal-Wallis test followed by Dunn’s multiple comparisons test, or, given data were normal and homoscedastic, by one-way ANOVA followed by Tukey’s multiple comparisons test; * 0.01 ≤ *p* ≤ 0.05,** 0.001 ≤ *p* ≤ 0.01, *** *p* ≤ 0.001 c) Spearman’s correlograms between MRI signals: dotted lines for linear regressions, coloured areas for 95%-confidence intervals, *R* = Spearman correlation coefficient, *p* values from the Spearman’s rank correlation test. d) 3D reconstructions of example *aLPC* and, respectively, e) *aCTRL* lesions. Note that T2FLAIR hyperintense signals take up the largest volume, followed by T1Gd and, depending on the lesion localization, DIR signals, while the SWI signals are mostly confined to the needle trajectory through the tissue.

As a potential correlate for demyelination, LPC-induced lesions exhibited significantly larger T2FLAIR hyperintensity than their age-matched control counterparts at the acute stage (T2FLAIR mean ± s.e.m.: *aLPC* = 535.1 ± 188.9 mm³ vs. *aCTRL* = 93.7 ± 63 mm³; Kruskal-Wallis test followed by Dunn’s multiple comparisons test *H*(1) = 8.92, *p* = 0.003; Fig. 3b). This was corroborated by a significantly higher volume of Gd uptake in T1 sequences in LPC lesions than in CTRL, suggestive of blood-brain-barrier breakdown in the acute stage (T1Gd mean ± s.e.m.: *aLPC* = 390.3 ± 96.7 mm³ vs. *aCTRL* = 31.7 ± 13.2 mm³; Kruskal-Wallis test followed by Dunn’s multiple comparisons test *H*(1) = 11.84, *p* < 0.001; Fig. 3b). In instances involving WM lesions with sufficient proximity to the cortex to potentially elicit juxtacortical demyelination, we examined the lesion susceptibility in DIR sequences: the LPC-induced lesions demonstrated significantly higher DIR hyperintensity than their control counterparts (DIR mean ± s.e.m.: *aLPC* = 227.2 ± 94.7 mm³ vs. *aCTRL* = 12.1 ± 5.6 mm³; Kruskal-Wallis test followed by Dunn’s multiple comparisons test *H*(1) = 11.31, *p* < 0.001; Fig. 3b). We included in this context a three-dimensional (3D) magnetization-prepared rapid acquisition with gradient echo (MPRAGE) sequence, a technique useful for confirming the exact localization (e.g. juxtacortical vs cortical vs mixed) of grey matter lesions in MS due to improved spatial resolution and signal-to-noise ratio compared to DIR(45) and confirmed significantly larger hypointense signals in the LPC study arm, 3D-MPRAGE mean ± s.e.m.: *aLPC* = 78.9 ± 46.0 mm³ vs. *aCTRL* = 3.9 ± 2.0 mm³; Kruskal-Wallis test followed by Dunn’s multiple comparisons test *H*(1) = 4.66, *p* = 0.031 (Fig. 3b). However, we noticed no difference with respect to the volume of SWI susceptibility between the intervention and the control arm, SWI mean ± s.e.m.: *aLPC* = 48.6 ± 7.8 mm³ vs. *aCTRL* = 51.1 ± 7.6 mm³; one-way ANOVA followed by Tukey’s multiple comparisons test *F*(1,25) = 0.11, *p* = 0.74 (Fig. 3b).

We observed recovery of the signal changes in the acute stage across the following 14-21 days, suggesting restorative tissue changes that could include remyelination, as is the case in, for instance,d rodent LPC models(17). While the degree of T2FLAIR hyperintensity became similar in the LPC group (T2FLAIR mean ± s.e.m., *iLPC* = 189.0 ± 116.5 mm³, a 65%-decrease from *aLPC* vs. *iCTRL* = 30.7 ± 10.5 mm³, a 67%-decrease from *aCTRL*; Kruskal-Wallis test followed by Dunn’s multiple comparisons test *H*(1) = 3.13, *p* = 0.07; Fig. 3b), there was still slightly higher T1Gd uptake in the LPC group at the intermediate stage (T1Gd mean ± s.e.m.: *iLPC* = 152.0 ± 90.0 mm³, a 61%-decrease from *aLPC* vs. *iCTRL* = 6.1 ± 2.3 mm³, an 81%-decrease from *aCTRL*; Kruskal-Wallis test followed by Dunn’s multiple comparisons test *H*(1) = 4.87, *p* = 0.03; Fig. 3b). In none of the other consensus MRI sequences, neither at the intermediate, nor at the subacute stage, did we observe any significant signal differences between the treatment and control arm, with a tendency towards attenuation of differences between the treatment arms in the subacute, compared to the intermediate stage, especially concerning T2FLAIR (T2FLAIR mean ± s.e.m.: *sLPC* = 46.4 ± 44.4 mm³ vs. *sCTRL* = 2.0 ± 2.0 mm³; Kruskal-Wallis test followed by Dunn’s multiple comparisons test *H*(1) = 0.48, *p* = 0.49; Fig. 3b) and T1Gd uptake volumes (T1Gd mean ± s.e.m.: *sLPC* = 38.4 ± 4.2 mm³ vs. s*CTRL* = 15.1 ± 15.1 mm³; Kruskal-Wallis test followed by Dunn’s multiple comparisons test *H*(1) = 1.21, *p* = 0.27; Fig. 3b).

To reinforce the hypothesis that the LPC-induced tissue effect, compatible with, but not as yet demonstrated to be a demyelination, drove the MRI signal values in the treatment group at the acute stage, we evaluated the correlation between the T2FLAIR and T1Gd against the correlation between T2FLAIR and SWI signals. We observed a strong positive correlation between T2FLAIR and T1Gd signal volumes in the acute stage in the treatment group (Spearman’s *R* = 0.8, *p* < 0.001), but no correlation in the control group (Spearman’s *R* = 0.34, *p* = 0.301; Fig. 3c). This could be reproduced when analysing cortical lesions: we found again a strong positive correlation between T2FLAIR and DIR signal volumes at the acute stage in the treatment (Spearman’s *R* = 0.87, *p* < 0.001; Fig. 3c), but not in the control group (Spearman’s *R* = 0.45, *p* = 0.162; Fig. 3b). However, when analysing T2FLAIR against SWI signal volumes at the acute stage, we found no correlation in the treatment (Spearman’s *R* = 0.29, *p* = 0.273; Fig. 3c) or in the control group (Spearman’s *R* = 0.55, *p* = 0.082; Fig. 3c).

These trends could be reproduced, albeit more attenuated, at the intermediate stage: T2FLAIR and T1Gd volumes showed a positive correlation in the treatment (Spearman’s *R* = 0.64, *p* = 0.034), but not in the control group (Spearman’s *R* = 0.46, *p* = 0.214), whereas T2FLAIR and SWI volumes exhibited no correlation in the treatment (Spearman’s *R* = 0.4, *p* = 0.226) or in the control group (Spearman’s *R* = 0.64, *p* = 0.064) (Suppl. Fig. 2). At the subacute stage, we observed no relevant correlations between the signal volumes, not even in the treatment group, between T2FLAIR and T1Gd (Spearman’s *R* = 0.2, *p* = 0.917) (Suppl. Fig. 2).

### *In vivo* PET-MRI and *ex vivo* histological analysis

We traced back tissue section locations by comparing anatomical landmarks in the histological slices to the coronal reconstruction of the MRI dataset (Fig. 4a). We ensured nearly spherical LPC/PBS-diffusion profiles of at least 5 mm radius via CED, in order to satisfy the minimum resolution requirements of the PET scanner.

**Figure 4.**
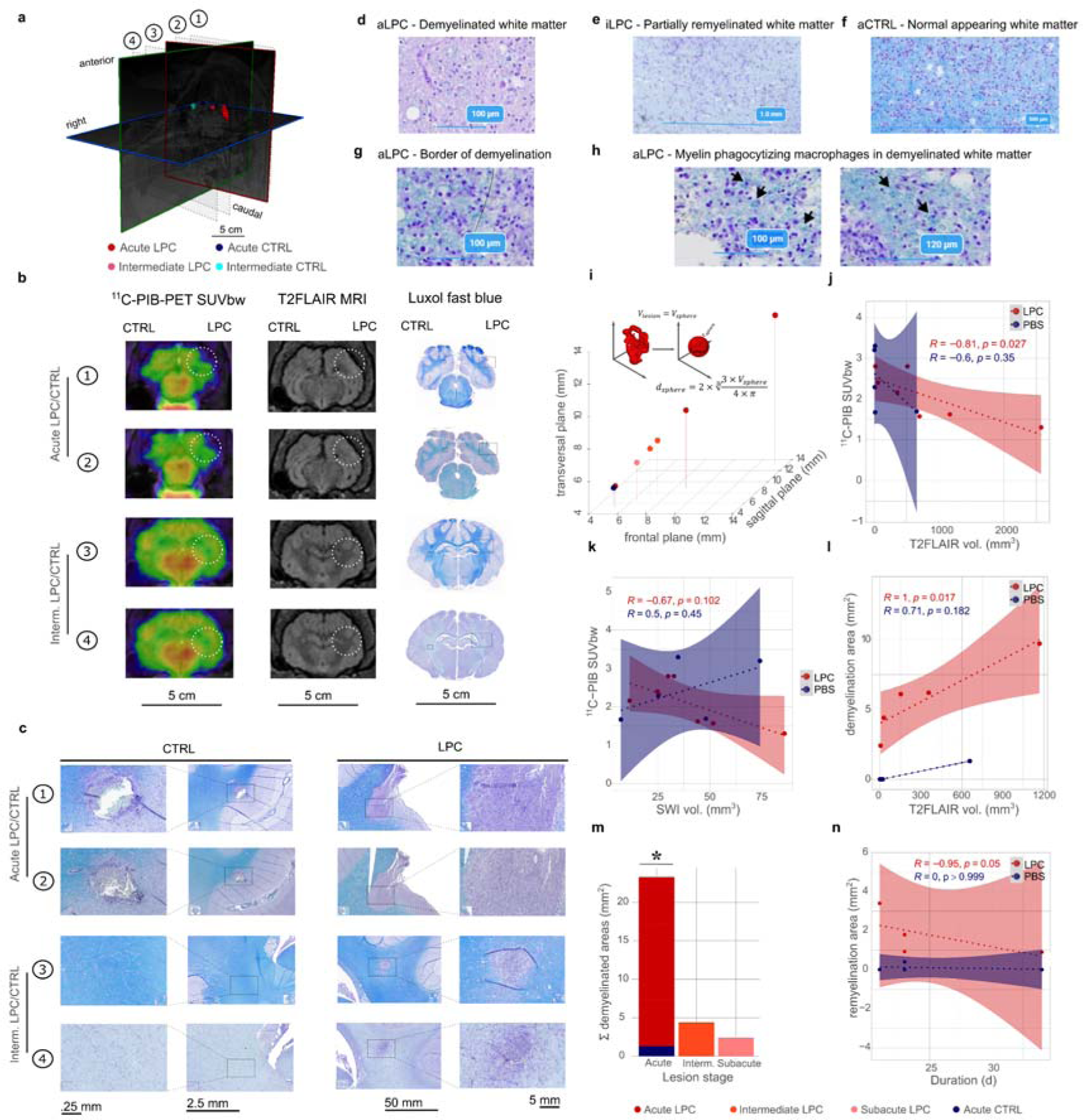
Multimodal validation of de- and remyelination. a) Example 3D lesion reconstruction. Dotted planes correspond to localizations in the following: 1 – *aLPC*, 2 – *aCTRL*, 3 – *iLPC,* 4 – *iCTRL*. b) Correlative imaging and histopathology. Dotted white circles on PET/MRI correspond to black dotted rectangles in LFB photomicrographs. c) Higher magnification photomicrographs (note different scales, for presentation purposes, because LPC lesions are of an order of magnitude more extensive than CTRL) corresponding to the aforementioned areas. d) High magnification micrograph of LFB-scarce WM in *aLPC*. e) Partially, diffusely remyelinated WM in *iLPC*, corresponding to a region from c), rows 3-4. f) Normal appearing WM in *aCTRL*. g) Well delineated (black line) border of demyelination in *aLPC*. h) Additional indicators of fresh demyelination in the form of myelin phagocytizing macrophages (foam cells) in *aLPC*. i) Equivolumetric sphere size distribution for *LPC* calculated by the depicted formula (*V* = volume, *d* = diameter). j) Spearman’s correlogram between the T2FLAIR volume and ^11^C-PIB-uptake in *aLPC/aCTRL*. k) Spearman’s correlogram between SWI susceptibility volumes and ^11^C-PIB-uptake in *aLPC/aCTRL*. l) Spearman’s correlogram between T2FLAIR volume and the histopathologically detectable mean demyelination areas in *aLPC/aCTRL*. m) Slide-by-slide sum (Σ) of demyelinated areas across groups. Statistical significance and *p* values determined by Kruskal-Wallis test followed by Dunn’s multiple comparisons test, * 0.01 ≤ *p* ≤ 0.05. n) Spearman’s correlogram between time post-induction and mean remyelination area. *R* = Spearman correlation coefficient, *p* values are calculated from Spearman’s rank correlation test.

Section-by-section analysis of ^11^C-PIB-PET and T2FLAIR-MRI *in vivo* tomographic sections and Luxol fast blue-stained and haematoxylin-and-eosin *ex vivo* tissue sections allowed to pinpoint the corresponding lesion locations across multiple modalities and timepoints (Fig. 4b and Fig. 4c, Suppl. Fig. 3). Demyelination was confirmed not only by diminished Luxol fast blue uptake (Fig. 4d-g), but also by myelin inclusions in PAS-stained macrophages(46) (Fig. 4h).

For every lesion we calculated the volume after segmentation in the T2FLAIR sequences and derived from this volume the diameter of a sphere with equal volume to the initial lesion. Accordingly, all lesions across the acute stage from both the treatment and the control group fulfilling all other inclusion criteria (no haemorrhagic component of more than 100mm³) exhibited an equivalent sphere diameter of above 5 mm (Fig. 4i). There was an inverse correlation between the T2FLAIR hyperintensity and ^11^C-PIB standard-uptake-values corrected by weight (*SUVbw*) in the treatment (Spearman’s *R* = -0.81, *p* = 0.027), but not in the control group (Spearman’s *R* = -0.6, *p* = 0.35), signifying that at least part of the T2FLAIR hyperintense area exhibited a reduced myelin content (Fig. 4j). Conversely, we could not determine any correlation between SWI volume and ^11^C-PIB *SUVbw*, neither in the LPC (Spearman’s *R* = -0.67, *p* = 0.102), nor in the CTRL group (Spearman’s *R* = 0.5, *p* = 0.45) (Fig. 4k). Pooled analysis of lesions across all stages was justified by the fact that T2FLAIR sequences cannot distinguish between de- and remyelination. In order to support the argument that the T2FLAIR signal changes in the acute stage were indeed driven by demyelination processes, we demonstrated a positive correlation between T2FLAIR hyperintensity volumes and demyelinated surface in the LPC treatment (Spearman’s *R* = 1, *p* = 0.017), but not in the control group (Spearman’s *R* = 0.71, *p* = 0.182) (Fig. 4l).

Neuropathological analysis of sections by slice-by-slice summation of segmented areas from both groups (derived from the rectangular method of morphometric estimation(34, 35), see Methods) revealed indeed a significantly higher cumulative demyelinated area in the acute treatment group than in the control group (demyelinated area mean ± s.e.m.: *LPC* = 7.2 ± 0.9 mm², n = 4, vs. *CTRL* = 0.3 ± 0.3 mm², n = 4; Kruskal-Wallis test *H*(1) = 5.6, *p* = 0.018). Conversely, partial remyelination was observed in the treatment group, but not in the control group (remyelinated area mean ± s.e.m.: *LPC* = 1.8 ± 0.6 mm², n = 4, vs. *CTRL* = 0.1 ± 0.1 mm², n = 4; Kruskal-Wallis test *H*(1) = 5.6, *p* = 0.018; Fig. 4m). Remyelination rate was highest during the earlier stages (around 14 days post-induction) and subsided towards the subacute stage (inverse correlation between remyelination surface and duration after lesion induction in the treatment, but not in the control group, Spearman’s *R* = -0.95, *p* = 0.051, Fig. 4n).

### Immunohistological and immunofluorescent characterization

Although LFB provides specific myelin staining of brain tissue and can be used to reliably track myelination status(47), the simple light-microscopic quantification of LFB across tissue surface does not take into account the other cellular components of a lesion, such as oligodendrocytes, astrocytes, neuronal somata/axons, microglia/macrophages and lymphocytes, which are relevant for determining how long demyelination is active (infiltration by lymphocytes and microglia/macrophages, presence of foamy cells), whether demyelination occurs along with oligodendrocyte loss, and, if so, whether there is oligodendrocytic repopulation during remyelination, whether axon pathology is present (as in a potential secondary neurodegeneration), and whether remyelination is restorative or only partial, but coupled with ongoing chronic and potentially neurotoxic inflammation and concurrent astrogliosis.

To answer these questions, we performed immunostaining of markers of relevant cellular types across all lesion stages: GFAP (for astrocytes), Iba1 (for microglia), Olig2 (for oligodendrocytes), NeuN (for neuronal somata), CD3 (for lymphocytes) and CD68 (for macrophages), as well as double immunofluorescence for GFAP against MOG (astrocytes versus myelin) and, respectively, for NFH against MOG (axons versus myelin).

LFB clearly demonstrated an area of focal demyelination in *aLPC* at the site of injection, significantly exceeding that of the control group, *aCTRL* (Fig. 5a, Suppl. Fig. 4a). This was paralleled by dense cellular infiltration in *aLPC*, consisting of CD3+ lymphocytes, GFAP+ reactive astrocytes, distinguished by hypertrophy and increased numbers of astrocytic branches and end feet, and Iba1+ activated microglia, distinguished by hypertrophic cell bodies, amoeboid shape and thickened cellular processes, but overall few CD68+ monocytes/macrophages (Fig. 5a). Among the latter however, we could distinguish myelin-phagocytizing, so-called foamy cells (Fig. 4h). The burgeoning inflammatory and reactive macro-/microglial infiltration went hand-in-hand with dramatically decreased Olig2+ cell numbers in *aLPC* (Fig. 5a), but not in *aCTRL* (Suppl. Fig. 4a) as well as focally increased GFAP-fluorescence concomitant with reduced MOG-fluorescence in *aLPC* (Suppl. Fig. 5a), but not in *aCTRL* (Suppl. Fig. 6a). Reactive astrogliosis and activated microglia were still present in *iLPC* as opposed to *iCTRL*, albeit to a reduced extent compared to *aLPC* (Fig. 5b, Suppl. Fig. 4b). In *iLPC*, we observed a higher number of CD68+ infiltrating cells than in *aLPC*, suggesting that in contrast to rodents, where remyelination is largely completed around 14-21 dpi(48, 49), in the minipig even at 30 dpi there is coexistence between a subsiding, but still active demyelinating component and tissue restorative mechanisms.

**Figure 5.**
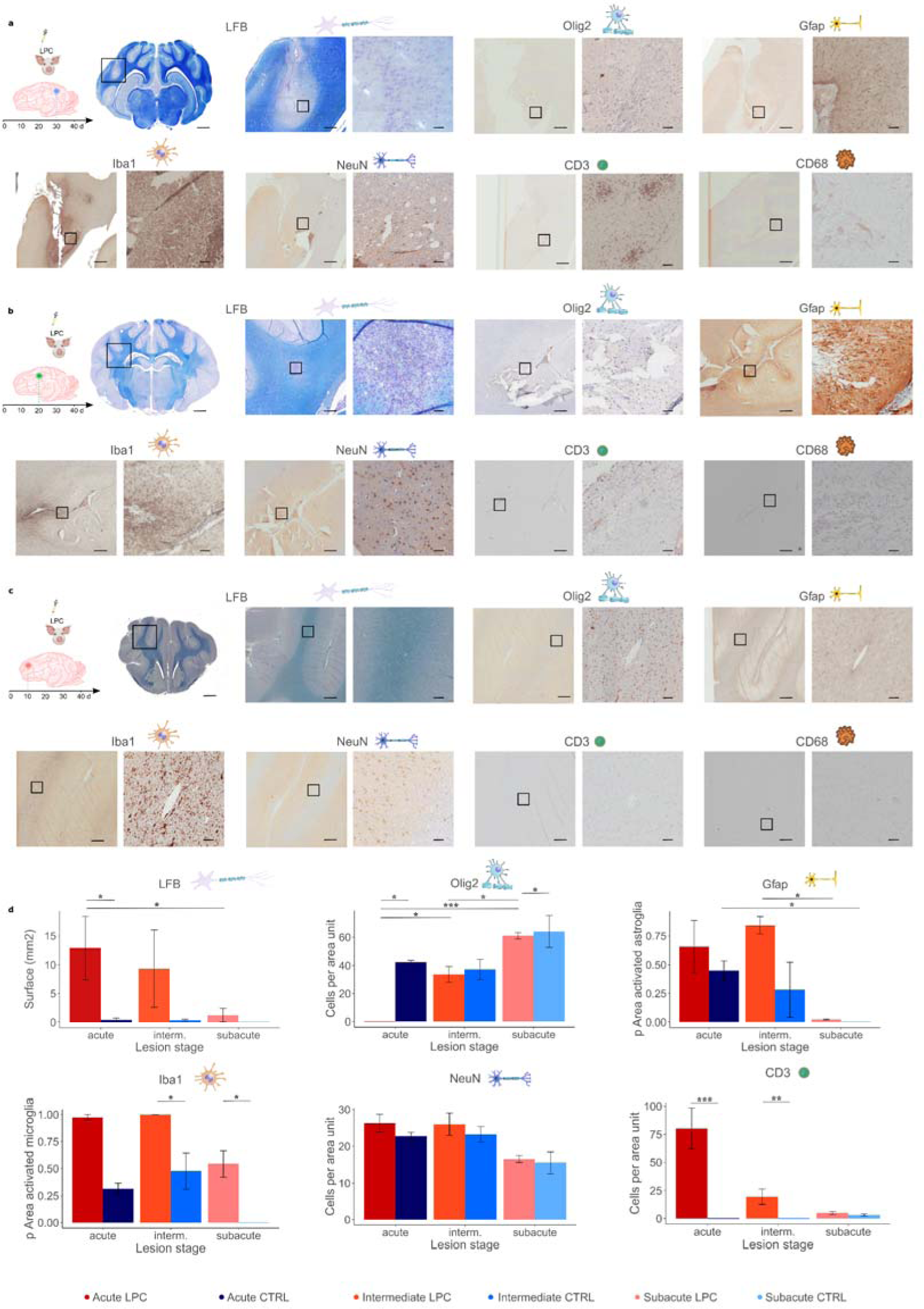
Immunohistochemical characterization of LPC-induced lesions across stages. a) Acute stage (*aLPC*), b) Intermediate (*iLPC*), c) Subacute (*sLPC*) Common denominators of a)-c): From left to right: Schematic corresponding to Fig. 1 of the lesion stage; LFB (Luxol fast blue) overview of entire coronal slice (scale bar = 5 mm, black square delineates ROI magnified on the right and in each low magnification inset of the following stainings). Cell-specific marker stainings containing paired low-magnification (15×, left, scale bar = 1 mm, black square delineates ROI magnified on the right) and high magnification insets (right, 60×, scale bar = 100 µm) for following cellular markers: Olig2 (oligodendrocytes), Iba1 (microglia), NeuN (neuronal somata), CD3 (lymphocytes), CD68 (monocytes/macrophages). d) From left to right: areas of LFB signal loss (i.e., demyelination), Olig2+ cell numbers per 300 × 300 µm² ROI, proportion of area containing activated GFAP+ astrocytes in 1 × 1 mm² ROI, proportion of area containing reactive Iba1+ microglia in 1 × 1 mm² ROI, NeuN+ cell numbers per 300 × 300 µm² ROI, CD3+ cell numbers per 300 × 300 µm² ROI. We refrained from CD68 quantification because of very low signal-to-noise ratio, presumably due to low cross-species primary antibody affinity. Statistical tests performed using Kruskal-Wallis-Test followed by Dunn’s multiple comparisons test. * 0.01 ≤ *p* ≤ 0.05, ** 0.01 ≤ *p* ≤ 0.001, *** *p* ≤ 0.001.

LFB reactivity as well as MOG-immunofluorescence were paralleled by the numbers of Olig2+ cells in the vicinity of the injection focus (Fig. 5a-c, Suppl. Fig. 5a-c), suggesting that demyelination occurred along with oligodendrocyte loss, while remyelination correlated with restoration of oligodendrocyte numbers. The latter, however, did not attain initial levels in *sLPC* when compared to *sCTRL* (Fig. 5d). Whether this had been paralleled by a lower content of myelin remained to be decided by the ultrastructural analysis (see below).

Parallel immunostaining for MOG and NfH revealed disrupted regularity in the distribution of neurofilaments (Suppl. Fig. 5a-c and Suppl. Fig. 6a-c) in all treatment groups, when compared to their respective controls. We did not find reduced numbers of neuronal somata in the cortical regions adjacent to the WM where the injections were placed. However, exact correspondence between the adjacent cortical areas where neuronal somata were counted and the precise axons contained in the WM tracts affected by the lesions is not a given in this setting, so that we resorted to clarify this question ultrastructurally as well (see below).

### Validation via electron-microscopy and against human data

As a further step towards validating the de- and remyelination surmised from the PET-MRI data and supported by conventional neuropathological studies, we performed scanning electron microscopy (SEM) from both the LPC and CTRL groups at either an acute or at a subacute timepoint (*aLPC*, *aCTRL*, *sLPC*, *sCTRL*). Moreover, we corroborated this with SEM biopsy data from two patients diagnosed and treated in our neurology department, Patient A (*aHum*) with an acute MS lesion in a demyelinating stage, and Patient S (*sHum*) with a subacute ADEM lesion in a remyelinating stage. However, due to the rarity of such biopsies even in a regional tertiary care centre such as ours, we were constrained to analysing WM samples from different human brain regions: *aHum* was taken from the cerebellar WM, while *sHum* was from the subcortical WM.

The acute as well as the subacute minipig control WM samples (*aCTRL, sCTRL*) featured an abundance of axonal fibres with intact myelin sheaths (Suppl. Fig. 7). In *aLPC* as well as in Patient A (*aHum*), however, we found that the axons exhibited features of pathological myelin: the myelin sheath was very thin, and we noticed myelin fragments in the extracellular space or intracellularly within lysosomal inclusions of phagocytizing cells (i.e., foamy cells), as well as, especially in *aLPC,* phagocytes engulfing myelin sheaths (Fig. 6a). The latter two features were not present anymore in *sLPC* or in Patient S (*sHum*), their place being taken by thinly, incompletely remyelinated axons (Fig. 6a). One difference we noted between *aLPC* and *aHum* pertained to the intracellular lipid droplets, which in *aLPC* appeared localized within phagocytes (macrophages or microglia), while in *aHum* they were more frequent within astrocytes (Fig. 6a). Thus, axonal diameter, as measured at the lesser axis, progressed from lowest in the acute stage (axonal diameter mean ± s.e.m., *aLPC* = 0.115 ± 0.01 µm, n = 746; *aLPC* < *aCTRL*, Kruskal-Wallis test followed by Dunn’s multiple comparisons test *H*(1) = 32.90, *df* =5, *p* < 0.001), to intermediate values in the subacute LPC lesion (axonal diameter mean ± s.e.m. *sLPC* = 0.22 ± 0.01 µm, n = 239; *sLPC* < *sCTRL*, Kruskal-Wallis test followed by Dunn’s multiple comparisons test *H*(1) = 14.08, *df* =5, *p* < 0.001, *sLPC* < *aCTRL* Kruskal-Wallis test followed by Dunn’s multiple comparisons test *H*(1) = 12.34, *df* =5, *p* < 0.001) and exhibited higher values both in the acute control and subacute control groups (axonal diameter mean ± s.e.m.: *aCTRL* = 0.444 ± 0.01 µm, n = 426; *sCTRL* = 0.653 ± 0.02 µm, n = 239) (Fig. 6b). This matched reports in patients with acute and subacute MS lesions(50), as well as the ratio among the data from both our patients (Patient A mean ± s.e.m.: *aHum* = 0.160 ± 0.01 µm, n = 1095, vs. Patient S mean ± s.e.m.: *sHum* = 0.301 ± 0.01 µm, n = 519) (Fig. 6b). As a more specific measure for the degree of myelination of an axon, since it transcends variations due to axon calibre, we quantified the g-ratio at 10 nm-resolution, defined as the ratio between the axon diameter and the diameter of the axon plus the myelin sheath. Thus, a higher g-ratio indicates a thinner myelin sheath, as in ongoing demyelination or incipient remyelination. In the minipig as well as in patient *sHum*, the sampled regions stemmed from the subcortical white matter, while in *aHum*, it originated from cerebellar WM, where in adulthood slightly different values can be expected(51). The g-ratio was highest in the acute LPC lesion (g-ratio mean ± s.e.m.: *aLPC* = 0.799 ± 0.01, n = 746; *aLPC* > *aCTRL*), took an intermediate value in the subacute LPC lesion, potentially correlating to the evidence of thinly and incompletely myelinated axons (g-ratio mean ± s.e.m.: *sLPC* = 0.708 ± 0.01, n = 327; *sLPC* > *sCTRL*, *sLPC* > *aCTRL*), and exhibited normal values in the rest of the minipig groups (g-ratio mean ± s.e.m.: a*CTRL* = 0.637 ± 0.01, n = 426; *sCTRL* = 0.626 ± 0.01, n = 239). There was a similar relationship in the patient data as well (Patient A mean ± s.e.m.: *aHum* = 0.697 ± 0.01, n = 1095, vs. Patient S mean ± s.e.m., *sHum* = 0.612 ± 0.01, n = 519) (Fig. 6b).

**Figure 6.**
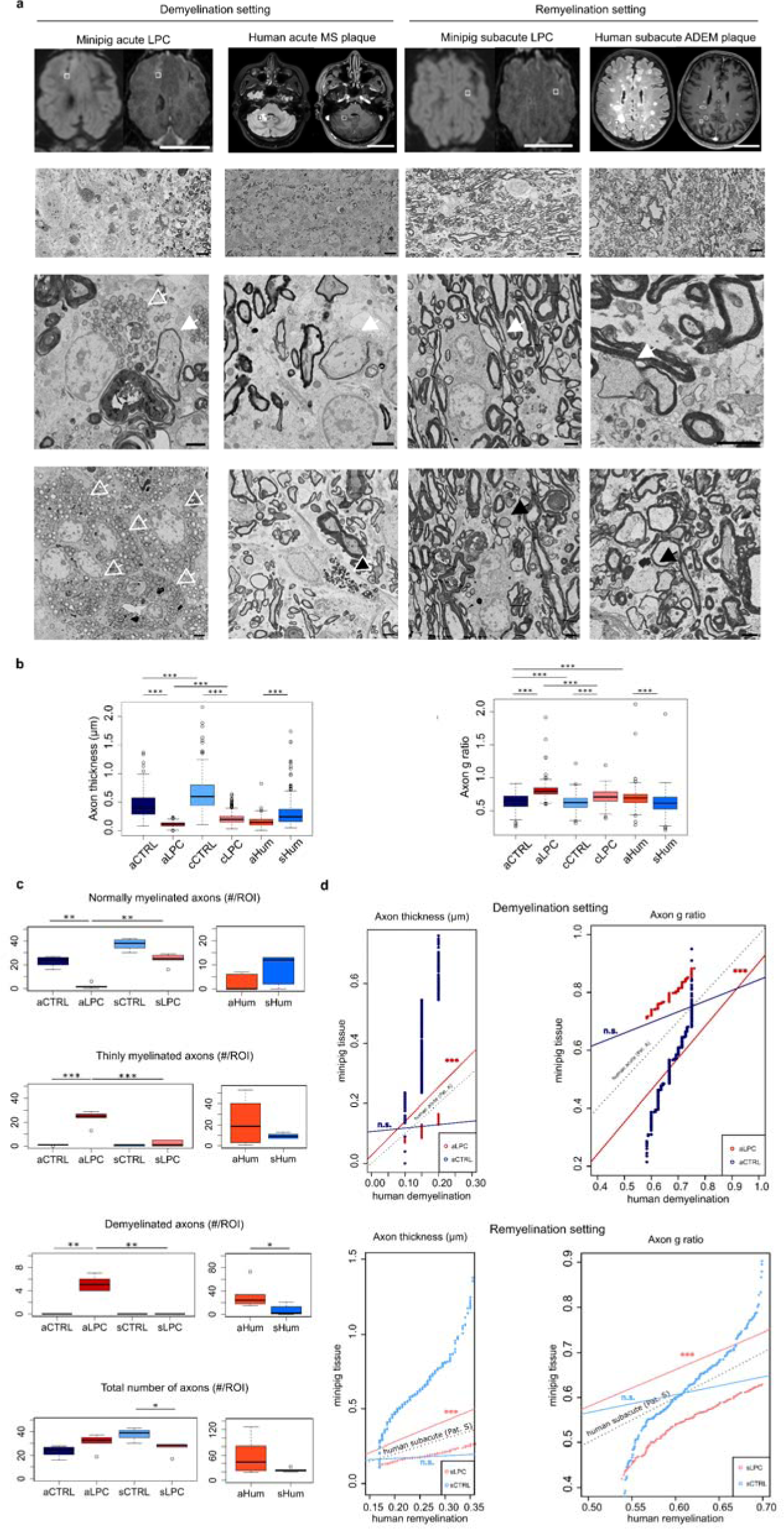
Comparison of MiniSWINE and human demyelinating CNS disease biopsy ultrastructure. a) 1^st^ row: Pairwise T2FLAIR and T1Gd axial MRI slices illustrating, in case of the minipigs (from the left, columns 1 and 3), the areas where autopsy was performed, and, in case of the humans (from the left, columns 2 and 4), the areas where biopsies were performed. White/black bounding boxes indicate approximate areas from which a small sample was analysed with SEM (see also Methods). 2^nd^ row: SEM-micrographs from *aLPC, aHum, sLPC* and, respectively, *sHum* groups, scale bar = 10 µm. 3^rd^ row, 4^th^ row: High magnification SEM-micrographs, scale bar = 2 µm. White arrows = thin myelin; hollow white arrows = lipid droplets in foam cells; black arrow with white outline: astrocytic lipid droplets and lysosomal inclusions; black arrows: axonal pathology. b) Quantification of axonal and myelin pathology. Mean ± s.e.m. of axonal diameter (above), as well as g-ratio (below) over all study groups, including *aHum* and *sHum*. c) Quantification of myelin pathology. Mean ± s.e.m. numbers of, from the left, normally myelinated, thinly myelinated and demyelinated axons per 10 × 10 µm ROI. b)-c) Kruskal-Wallis test followed by Dunn’s multiple comparisons test, * 0.01 ≤ *p* ≤ 0.05, ** 0.01 ≤ *p* ≤ 0.001, *** *p* ≤ 0.001. d) Comparisons of predictive power of linear regression models of axonal thickness and g-ratio from *MiniSWINE* (the linear model itself as a straight line) compared to the human linearized data, represented by the dotted straight line. The nearer the trajectory of the coloured line was to the one of the dotted line, the better the prediction of *MiniSWINE* relative to human data was. Statistical significance and *p* values were determined by likelihood ratio tests, *** *p* ≤ 0.001, n. s. = not significant.

We also consistently observed an increased proportion of pathological axons (partially myelinated or denudated) in *aLPC* and *sLPC*. In the former case, demyelinated axons were more frequent, while in the latter, thinly myelinated axons predominated (Fig. 6c). At the acute stage, the proportion of pathological axons over all axons was higher in *aLPC* (24% thinly myelinated, 5% demyelinated) than in *aCTRL* (8% thinly myelinated, 0% demyelinated), while in the subacute stage the proportion of remyelinated axons over all axons was higher in *sLPC* (2% thinly myelinated) than in *sCTRL* (0.6% thinly myelinated) (Fig. 6c). Overall, however, we found slightly lower numbers of axons (by around 15%, axon numbers per 10 × 10 µm² ROI, mean ± s.e.m.: *sLPC* = 25 ± 3.5, *sCTRL* = 28.83 ± 3.44, *sLPC* < *sCTRL*, Kurskal-Wallis p = 0.01) in *sLPC* (Fig. 6c), which, corroborated with data from NfH immunostaining, hinted to a certain degree of secondary axonal degeneration present at the subacute stage.

By adjusting linear models to the distributions of axonal diameter and g-ratios from minipig and human tissue and performing likelihood ratio tests (LR), we could further corroborate the modelling of *aHum* data by *aLPC* regarding both axonal diameter (*aLPC* significantly improving LR over *aCTRL*, *Df* = 3, *H²* = 263.83, *p* < 0.0001) as well as g-ratio (*aLPC* significantly improving LR over *aCTRL*, *Df* = 3, *H²* = 676.67, *p* < 0.0001). Analogous results were obtained for approximating *sHum* by *sLPC* vs *sCTRL* both regarding the axonal diameter and the g-ratio (Fig. 6c-d). Thus, minipig LPC tissue exhibited similarities in key features of de- and remyelination stages to human inflammatory demyelinating CNS disease tissue with regards to ultrastructurally-verifiable parameters. To further evaluate the feasibility of modelling human chronic-inflammatory demyelinating disease by *MiniSWINE*, we took advantage of the Kullback-Leibler Divergence (DKL) as a measure for information loss (measured in bits) when modelling one probability distribution (i.e., human acutely demyelinated CNS tissue axons) with another, simpler or more easily attainable data probability distribution (i.e., data from *MiniSWINE*). To this end, we compared the DKL when modelling the distributions of axonal thickness and g-ratio of, on the one hand, the human de- and, respectively, remyelinated tissue and, on the other hand, *aLPC/aCTRL* vs, respectively, *sLPC/sCTRL* minipig tissue. Regarding axonal thickness, the intrinsic discrepancy of approximating the *aHum* distribution was lower when using *aLPC* (DKL_aLPC = 0.0087 bits) than when using *aCTRL* tissue (DKL_aCTRL = 0.114 bits). This was reflected with respect to the g-ratio as well, where the intrinsic discrepancy was lower when using *aLPC* (DKL_aLPC = 0.0003 bits) than when using *aCTRL* (DKL_aCTRL = 0.019 bits). In case of the subacute, remyelinated setting (*sHum*), the intrinsic discrepancy regarding the axonal thickness when modelling via *sLPC* (DKL_sLPC = 0.0004 bits) also considerably smaller then when employing *sCTRL* (DKL_sCTRL = 0.043 bits). The g-ratio could also be approximated with less information loss when calculating via *sLPC* (DKL_sLPC = 0.0007 bits) than via *sCTRL* (DKL_sCTRL = 0.0082 bits).

## Discussion

We aimed in this study to demonstrate the feasibility of validly simulating acute demyelination followed by at least remyelination in a large animal model, reflecting what is known to occur in human inflammatory disorders of the CNS and allowing follow-up and characterization using multimodal imaging and microscopy. We chose minipigs to study de- and remyelination, (1) because they have relatively large gyrencephalic brains, (2) a high degree of neuroanatomical similarity to humans, including a similar white-to-grey matter (WM:GM) ratio (∼ 60:40), as well as (3) a manageable adult mass of around 60kg(9, 52). The latter aspect is relevant to stereotaxy compared to fattening pigs, since navigational coordinates for stereotaxy can shift during brain and body growth. This restricts longitudinal precision neuroanatomy studies to adult animals, which for fattening pigs, due to their size and body mass, would pose significant infrastructural problems in using analogous diagnostic devices as for human clinical use. Finally, minipigs are easily maintained in controlled conditions(9). For this, we induced reversible, spatially and temporally controlled demyelination using precise, stereotactic delivery of LPC via CED. To this end, we chose to develop an EMTS and a surgical-imaging-microscopy platform, *MiniSWINE*, which due to its modular construction is compatible with adaptations, e.g., to implement translational developments. One of the initial challenges was to design a cannula achieving an optimal, spherical substance spread function at canonical CED injection rates, which would also be compatible with readily available single-use catheters. In contrast to previously reported OTS, the EMTS we developed for this application was not limited by catheter, sensor wire or cannula bending in the tissue, since the tracking targeted the tip of the instrument and not the initial part of the instrument shaft. Also, as EMTS does not require visual access for a camera, the handling of the navigated drill guide by the operator on challengingly small surfaces such as the forehead is easier than in the case of OTS, by preventing line-of-sight issues faced by contemporary neurosurgical navigation systems.

However, a notable limitation of the *MiniSWINE* EMTS was interference with metals in the immediate vicinity of the surgical field, which could not be eliminated. Indeed, in one of forty instances we were confronted with magnetization of the cannula which required its replacement during surgery, an issue not occurring in OTS. This could be compensated for practical purposes by ensuring appropriate spacing between the EMTS and metallic surgical instruments, in a way that provided a satisfactory precision of injection. This, as well as the fact that salient anatomical points *in vivo*, as opposed to *in vitro*, are registered on skin, which can physiologically shift within certain limits in relation to subcutaneous tissue, contributed to the discrepancy between the navigation precision *in vitro* (mean ± s.e.m.: 0.23 ± 0.03 mm) versus *in vivo* (mean ± s.e.m.: 1.54 ± 0.4 mm).

An important consideration once having established the platform *MiniSWINE* from the technical standpoint, was validating it against the MAGNIMS consortium multimodal MRI sequences performed on the same devices in a clinical setting as would be the case for the diagnosis and follow-up of a chronic inflammatory demyelinating disorder in human patients, such as for MS. Intercorrelating, reproducibly higher T2FLAIR, T1Gd and DIR signal susceptibilities in the treatment, but not in the control group, indicated a tissue-effect, compatible with a demyelination, as well as a partial permeability increase of the blood-brain barrier (Gd uptake) triggered at the acute stage by the intraparenchymal LPC injection. The absence of correlations between SWI and the aforementioned sequences in the treatment as well as in the control group suggested that the inherent microhaemorrhage along the injection trajectory did not significantly confound LPC-induced effects. One interesting aspect of SWI-related MRI-sequences, which we did not investigate, in light of the known accumulation of iron in macrophages in active lesions and in shadow plaques(53), would be potential sensitivity to ongoing inflammation or even, in the case of quantitative susceptibility imaging (QSM), sensitivity to myelin content, which could be exploited as a biomarker for remyelinated lesions(54). Results from our longitudinal MRI follow-up alone were in principle compatible with subsequent remyelination, but since none of the current MAGNIMS consensus sequences can reliably track remyelination(2), we needed additional, more specific imaging as well as parallel histopathological characterization of lesion evolution to validate our hypothesis. As a proof-of-concept for testing minimally invasive, novel imaging biomarkers, that could help tackle the current challenge of detecting incipient remyelination, we resorted to *in vivo* [^11^C]-PIB-PET imaging, with the aid of which we could demonstrate a correlation between subsiding T2FLAIR and T1Gd signals and restoration of PIB-uptake, as a potential starting point for evaluating this technique in future (human) studies. We also showed via parallel investigation *in vivo* by MRI and *ex vivo* by histopathology that we could follow-up signals of demyelination for weeks without confounding due to potential intercurrent mechanical tissue damage or perilesional intracerebral haemorrhage.

The de-/remyelination dynamics observed in *MiniSWINE* differed to a certain extent from those described in rodent models(17, 55). The latency to remyelination took intermediate values between the 2-3 weeks described in rodent studies(17, 49) and the incomplete remyelination even 6 weeks post-induction in other large animal models(15, 16). This is in line with recent reports describing not only the lesion volume, but also species-inherent differences in OPC infiltration of the lesion, proliferation and differentiation as the main factors determining the remyelination status(16). We cannot exclude age-dependent effects on the latency of the remyelination(15), but consider this unlikely due to the homogenous cohort of young adult minipigs used (aged 19 ± 1 months, mean ± s.e.m., considering a life expectancy of 15-20 years). Future work with increased imaging density and specific labelling of OPCs may help further clarify the interspecies differences. We demonstrated experimentally that ultrastructural indicators of de- and remyelination from *MiniSWINE* matched human biopsy data from two patients in distinct phases of a demyelinating CNS inflammatory process, by exhibiting similar lesion characteristics. Nevertheless, we had to compare WM axonal diameters and g-ratios from different human brain regions, which restricted the comparability of results between the demyelinating and remyelinating human conditions. We observed homogenous de- and remyelination patterns irrespective of the injection location within the centrum semiovale, even though existing reports in the literature hint towards subtle interregional differences within different areas of the corpus callosum in rodent cuprizone models(49, 56).

Interestingly, we found a certain degree of putatively secondary axonal pathology (around 15% of axons) inherent to the subacute treatment group, which deviates from the paradigm of a pure demyelinating non-inflammatory LPC-induced lesion in rodent studies and more closely resembles results from rabbit and macaque studies(15, 16). This is to our knowledge the first description of subacute LPC-induced and altogether demyelinating lesion dynamics in the pig brain, since a previous study has focused solely on the acute stage(14). One potentially critical aspect here is that in the remyelination setting, the proportion of axons with thin myelin, typical for remyelinated axons, is quite low (on the order or 2%), which is counterintuitive, given that the lesions are still detectable by [^11^C]-PIB-PET-MRI. However, MRI sequences as stipulated by the MAGNIMS consortium do not specifically identify de- or remyelination events. Also, the [^11^C]-PIB uptake correlates with myelination, so that in a remyelinating lesion, one expects a lower contrast between the potential lesion and surrounding parenchyma, further lowering detectability. Therefore, astrogliosis, microglial and immune cell infiltration can also confound the imaging, especially in the case of the MRI, leading to an overestimation of the de-/remyelinated lesion core, which can, in our paradigm, be precisely determined on histological and ultrastructural basis.

On the whole, another critical aspect of the study was given by the low experimental numbers concomitantly examined with multimodal diagnostic methods, especially in the subacute group. This was due, on the one hand, to longitudinal imaging being performed in a single animal, leading to more lesions in the acute stage being generated than subacute lesions could emerge. On the other hand, we were constrained by local ethical regulations to not only establishing the pipeline sequentially, but at the same time to also generating final data. Despite these limitations, we established controlled cerebral de- and remyelination in a large animal model and validate it in a multimodal imaging and microscopy setting against existing and emerging biomarkers. In this sense, *MiniSWINE* opens the avenue for immediate clinical translation to diagnostic procedures and new imaging biomarkers, while representing a gateway to the potential future testing of novel remyelinating agents. Furthermore, due to its modular construction, the EMTS conceived by us may enhance stereotactic substance delivery in human neurosurgery.

## Contributors

BH conceived the study. MA, GKT, VMB and JG performed surgical experiments. SB, JF and JR provided veterinarian support across the entire study. CS, SH, BL, EL and TL designed, fabricated and coordinated the EMTS during neurosurgery. IY and SN directed PET imaging. FLS, JS, KM, GKT, SSA and MA performed histopathology. JK and MM directed MRI scans. TM and MS implemented the SEM pipeline and performed SEM analysis. MA and GKT performed data analysis. MA, GKT, IY, FLS, JK verified underlying data. MA and BH wrote the paper with contributions from all coauthors. All authors read and approved the final version of the manuscript.

## Data sharing statement

The main data supporting the results of this study can be made available upon reasonable request from the corresponding author.

## Acknowledgements

MA, MS, TM and BH were supported by DFG under Germany’s Excellence Strategy within the framework of the Munich Cluster for Systems Neurology (EXC 2145 SyNergy, ID 390857198); MA, MS and TM were further supported by TRR 274/1 2020, 408885537 (projects B03 and Z01). We thank W. Weber for essential infrastructural support and C. Baumgartner for critical reading of the manuscript. We also thank all technical assistance staff from the departments of Nuclear Medicine, Neuropathology, Neurology, Veterinary Medicine and from the German Centre for Neurodegenerative Diseases for their essential support during this study. We thank the manufacturers at Ergosurg GmbH for fabricating the instruments for the EMTS.

## References

1. Dendrou CA, Fugger L, Friese MA. Immunopathology of multiple sclerosis. Nature Reviews Immunology. 2015;15:545–58.

2. Filippi M, Brück W, Chard DT, Fazekas F, Geurts JJG, Enzinger C, et al. Association between pathological and MRI findings in multiple sclerosis. The Lancet Neurology. 2019;18:198–210.

3. Plemel JR, Liu W-Q, Yong VW. Remyelination therapies: a new direction and challenge in multiple sclerosis. Nature Reviews Drug Discovery. 2017;16:617–34.

4. Walhovd KB, Johansen-Berg H, Káradóttir RT. Unraveling the secrets of white matter@ Bridging the gap between cellular, animal and human imaging studies. Neuroscience. 2014;276:2–13.

5. Ransohoff RM. Animal models of multiple sclerosis: the good, the bad and the bottom line. Nature Neuroscience. 2012;15:1074 - 7.

6. Procaccini C, De Rosa V, Pucino V, Formisano L, Matarese G. Animal models of Multiple Sclerosis. European Journal of Pharmacology. 2015;759:182–91.

7. Blakemore WF, Franklin RJM. Remyelination in experimental models of toxin-induced demyelination. Current topics in microbiology and immunology. 2008;318:193–212.

8. Fischer K, Schnieke A. Extensively edited pigs. Nature biomedical engineering. 2021;5 2:128–9.

9. Ardan T, Baxa M, Levinská Be, Sedlâ ková M, Nguyen TD, Klima J, et al. Transgenic minipig model of Huntington’s disease exhibiting gradually progressing neurodegeneration. Disease Models & Mechanisms. 2019;13.

10. Flisikowska T, Egli J, Flisikowski K, Stumbaum M, Küng E, Ebeling M, et al. A humanized minipig model for the toxicological testing of therapeutic recombinant antibodies. Nature biomedical engineering. 2022.

11. Wakeman DR, Crain AM, Snyder EY. Large animal models are critical for rationally advancing regenerative therapies. Regenerative medicine. 2006;1 4:405–13.

12. Hou N, Du X, Wu S. Advances in pig models of human diseases. Animal Model Exp Med. 2022;5(2):141–52.

13. Singer BA, Tresser NJ, Frank JA, McFarland HF, Biddison WE. Induction of experimental allergic encephalomyelitis in the NIH minipig. Journal of Neuroimmunology. 2000;105(1):7–19.

14. Kalkowski L, Malysz-Cymborska I, Golubczyk D, Janowski M, Holak P, Milewska K, et al. MRI-guided intracerebral convection-enhanced injection of gliotoxins to induce focal demyelination in swine. PLoS ONE. 2018;13.

15. Cooper JJM, Polanco JJ, Saraswat D, Peirick JJ, Seidl A, Li Y, et al. Chronic demyelination of rabbit lesions is attributable to failed oligodendrocyte progenitor cell repopulation. Glia. 2023;71(4):1018–35.

16. Sarrazin N, Chavret-Reculon E, Bachelin C, Felfli M, Arab R, Gilardeau S, et al. Failed remyelination of the nonhuman primate optic nerve leads to axon degeneration, retinal damages, and visual dysfunction. Proc Natl Acad Sci U S A. 2022;119(10):e2115973119.

17. Cantuti-Castelvetri L, Fitzner D, Bosch-Queralt M, Weil M-T, Su M, Sen P, et al. Defective cholesterol clearance limits remyelination in the aged central nervous system. Science. 2018;359:684–8.

18. Mozafari S, Deboux C, Laterza C, Ehrlich M, Kuhlmann T, Martino G, et al. Beneficial contribution of induced pluripotent stem cell-progeny to Connexin 47 dynamics during demyelination-remyelination. Glia. 2021;69(5):1094–109.

19. Hall SM. The effect of injections of lysophosphatidyl choline into white matter of the adult mouse spinal cord. J Cell Sci. 1972;10(2):535–46.

20. Denic A, Johnson AJ, Bieber AJ, Warrington AE, Rodriguez M, Pirko I. The relevance of animal models in multiple sclerosis research. Pathophysiology. 2011;18(1):21–9.

21. Koivukangas T, Katisko JPA, Koivukangas JP. Technical accuracy of optical and the electromagnetic tracking systems. SpringerPlus. 2013;2.

22. Pawlowsky K, Ernst L, Steitz J, Stopinski T, Kögel B, Henger A, et al. The Aachen Minipig: Phenotype, Genotype, Hematological and Biochemical Characterization, and Comparison to the Göttingen Minipig. European Surgical Research. 2017;58(5-6):193–203.

23. Bobo RH, Laske DW, Akbasak A, Morrison PF, Dedrick RL, Oldfield EH. Convection-enhanced delivery of macromolecules in the brain. Proc Natl Acad Sci U S A. 1994;91(6):2076–80.

24. Sillay KA, McClatchy SG, Shepherd BA, Venable GT, Fuehrer TS. Image-guided convection-enhanced delivery into agarose gel models of the brain. J Vis Exp. 2014(87).

25. Yamazaki R, Ohno N, Huang JK. Acute motor deficit and subsequent remyelination-associated recovery following internal capsule demyelination in mice. Journal of Neurochemistry. 2021;156(6):917–28.

26. Wattjes MP, Ciccarelli O, Reich DS, Banwell BL, Stefano ND, Enzinger C, et al. 2021 MAGNIMs CMSC NAIMS consensus recommendations on the use of MRI in patients with multiple sclerosis. The Lancet Neurology. 2021;20:653–70.

27. Brant-Zawadzki MN, Gillan GD, Nitz WR. MP RAGE: a three-dimensional, T1-weighted, gradient-echo sequence--initial experience in the brain. Radiology. 1992;182 3:769–75.

28. Riche M, Marijon P, Amelot A, Bielle F, Mokhtari K, Chambrun MP, et al. Severity, timeline, and management of complications after stereotactic brain biopsy. J Neurosurg. 2022;136(3):867–76.

29. Stankoff B, Freeman L, Aigrot M-S, Chardain A, Dollé F, Williams AC, et al. Imaging central nervous system myelin by positron emission tomography in multiple sclerosis using [methyŒ 11C] R (T2 methylaminophenylI V hydroxybenzothiazole. Annals of Neurology. 2011;69.

30. Mathis CA, Wang Y, Holt DP, Huang G-f, Debnath ML, Klunk WE. Synthesis and evaluation of 11C-labeled 6-substituted 2-arylbenzothiazoles as amyloid imaging agents. Journal of medicinal chemistry. 2003;46 13:2740–54.

31. Lopresti BJ, Klunk WE, Mathis CA, Hoge JA, Ziolko SK, Lu X, et al. Simplified quantification of Pittsburgh Compound B amyloid imaging PET studies: a comparative analysis. Journal of nuclear medicine : official publication, Society of Nuclear Medicine. 2005;46 12:1959–72.

32. Zhanmu O, Yang X, Gong H, Li X. Paraffin-embedding for large volume bio-tissue. Scientific Reports. 2020;10.

33. Schindelin J, Arganda-Carreras I, Frise E, Kaynig V, Longair M, Pietzsch T, et al. Fiji: an open-source platform for biological-image analysis. Nature Methods. 2012;9(7):676-82.

34. Rosen GD, Harry JD. Brain volume estimation from serial section measurements: a comparison of methodologies. J Neurosci Methods. 1990;35(2):115–24.

35. Gundersen HJ, Jensen EB. The efficiency of systematic sampling in stereology and its prediction. J Microsc. 1987;147(Pt 3):229–63.

36. Kislinger G, Gnägi H, Kerschensteiner M, Simons M, Misgeld T, Schifferer M. ATUM-FIB microscopy for targeting and multiscale imaging of rare events in mouse cortex. STAR Protocols. 2020;1(3):100232.

37. Stikov N, Campbell JSW, Stroh T, Lavelée M, Frey S, Novek J, et al. Quantitative analysis of the myelin g-ratio from electron microscopy images of the macaque corpus callosum. Data in Brief. 2015;4:368–73.

38. Holman C, Piper SK, Grittner U, Diamantaras A-A, Kimmelman J, Siegerink B, et al. Where Have All the Rodents Gone? The Effects of Attrition in Experimental Research on Cancer and Stroke. PLoS Biology. 2016;14.

39. NDI. Aurora V3.1 User Guide. 2018; Revision 4.

40. Pomfret RJ, Miranpuri GS, Sillay KA. The Substitute Brain and the Potential of the Gel Model. Annals of Neurosciences. 2013;20:118–22.

41. Miranpuri GS, Hinchman A, Wang A, Schomberg DT, Kubota K, Brady M, et al. Convection Enhanced Delivery: A Comparison of infusion characteristics in ex vivo and in vivo non-human primate brain tissue. Annals of Neurosciences. 2013;20:108–14.

42. Mehta AM, Sonabend AM, Bruce JN. Convection-Enhanced Delivery. Neurotherapeutics. 2017;14:358–71.

43. Ali S. joimax® Intracs® em System. In: Services DoHaH, editor. June 26, 2020.

44. Thompson AJ, Banwell BL, Barkhof F, Carroll WM, Coetzee T, Comi G, et al. Diagnosis of multiple sclerosis: 2017 revisions of the McDonald criteria. The Lancet Neurology. 2018;17:162–73.

45. Nelson F, Poonawalla AH, Hou P, Wolinsky JS, Narayana PA. 3D MPRAGE improves classification of cortical lesions in multiple sclerosis. Multiple Sclerosis. 2008;14:1214 - 9.

46. Huitema MJD, Strijbis EM, Luchicchi A, Bol JGJM, Plemel JR, Geurts JJG, et al. Myelin Quantification in White Matter Pathology of Progressive Multiple Sclerosis Post-Mortem Brain Samples: A New Approach for Quantifying Remyelination. International Journal of Molecular Sciences. 2021;22.

47. Kluver H, Barrera E. A method for the combined staining of cells and fibers in the nervous system. J Neuropathol Exp Neurol. 1953;12(4):400–3.

48. Cantuti-Castelvetri L, Fitzner D, Bosch-Queralt M, Weil MT, Su M, Sen P, et al. Defective cholesterol clearance limits remyelination in the aged central nervous system. Science. 2018;359(6376):684–8.

49. Werkman IL, Lentferink DH, Baron W. Macroglial diversity: white and grey areas and relevance to remyelination. Cellular and Molecular Life Sciences. 2021;78(1):143–71.

50. Albert M, Antel JP, Brück W, Stadelmann C. Extensive Cortical Remyelination in Patients with Chronic Multiple Sclerosis. Brain Pathology. 2007;17.

51. Bouhrara M, Kim RW, Khattar N, Qian W, Bergeron C, Melvin D, et al. Age-related estimates of aggregate g-ratio of white matter structures assessed using quantitative magnetic resonance neuroimaging. Human Brain Mapping. 2021;42:2362 - 73.

52. Vodj ka P, Smetana K, DvoYánková B, Emerick T, Xu Y, Ourednik J, et al. The Miniature Pig as an Animal Model in Biomedical Research. Annals of the New York Academy of Sciences. 2005;1049.

53. Popescu BF, Frischer JM, Webb SM, Tham M, Adiele RC, Robinson CA, et al. Pathogenic implications of distinct patterns of iron and zinc in chronic MS lesions. Acta Neuropathologica. 2017;134(1):45–64.

54. Rahmanzadeh R, Galbusera R, Lu PJ, Bahn E, Weigel M, Barakovic M, et al. A New Advanced MRI Biomarker for Remyelinated Lesions in Multiple Sclerosis. Ann Neurol. 2022;92(3):486–502.

55. Cunha MI, Su M, Cantuti-Castelvetri L, Müller SA, Schifferer M, Djannatian M, et al. Pro-inflammatory activation following demyelination is required for myelin clearance and oligodendrogenesis. The Journal of Experimental Medicine. 2020;217.

56. Steelman AJ, Thompson JP, Li J. Demyelination and remyelination in anatomically distinct regions of the corpus callosum following cuprizone intoxication. Neurosci Res. 2012;72(1):32–42.

