## Supplementary Material for "Multimodally trackable and clinically translatable platform for modelling human demyelinating brain diseases by temporally dispersed chemically induced lesions in the pig brain"

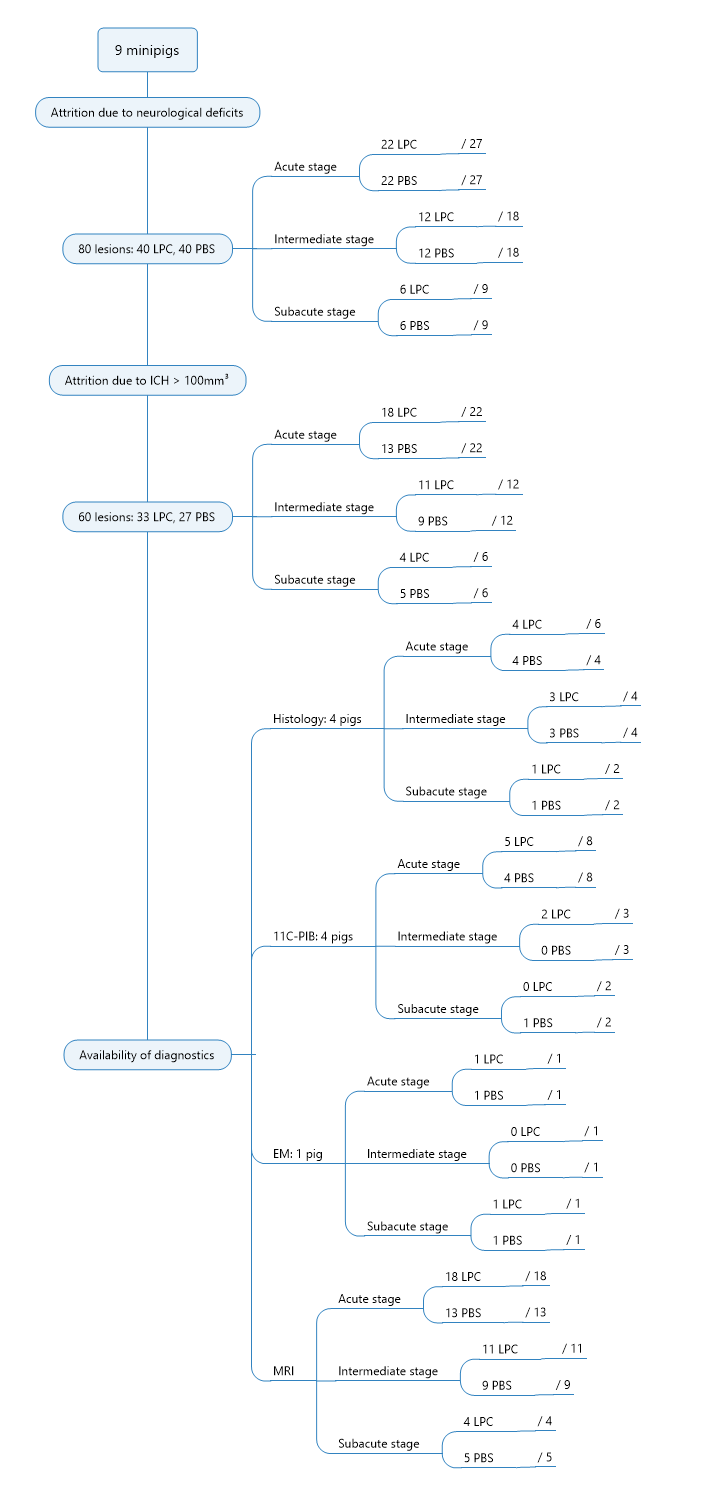

**Supplementary Figure 1.** *Attrition rates*. Overview of experimental numbers and limitations due to attrition because of neurological deficits, which constituted an ethical termination criterion, size of perilesional intracerebral haemorrhage (ICH) over 100 mm³ as well as the availability of diagnostic methods, which could not cover all minipigs involved in this study. Numbers, if not stated otherwise, refer to lesions as statistical units. The lower-level subtopics, introduced by “/”, refer to the maximum number of expectable samples after the previous step of attrition. LPC = treatment group, PBS = control group.

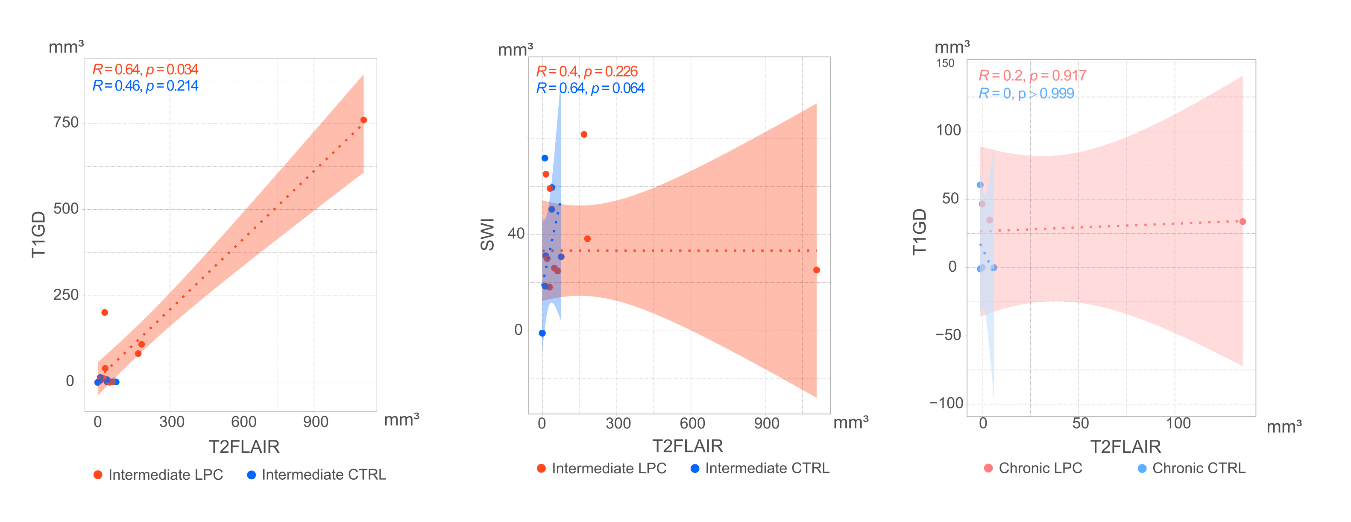

**Supplementary Figure 2.** *Intermediate and subacute stage MRI sequence correlations***.** Spearman’s correlograms between MRI signals: points represent individual values, dotted lines represent linear regression models, the coloured areas represent 95%-confidence intervals, R is the Spearman correlation coefficient, *p* values are calculated from the Spearman’s rank correlation test.

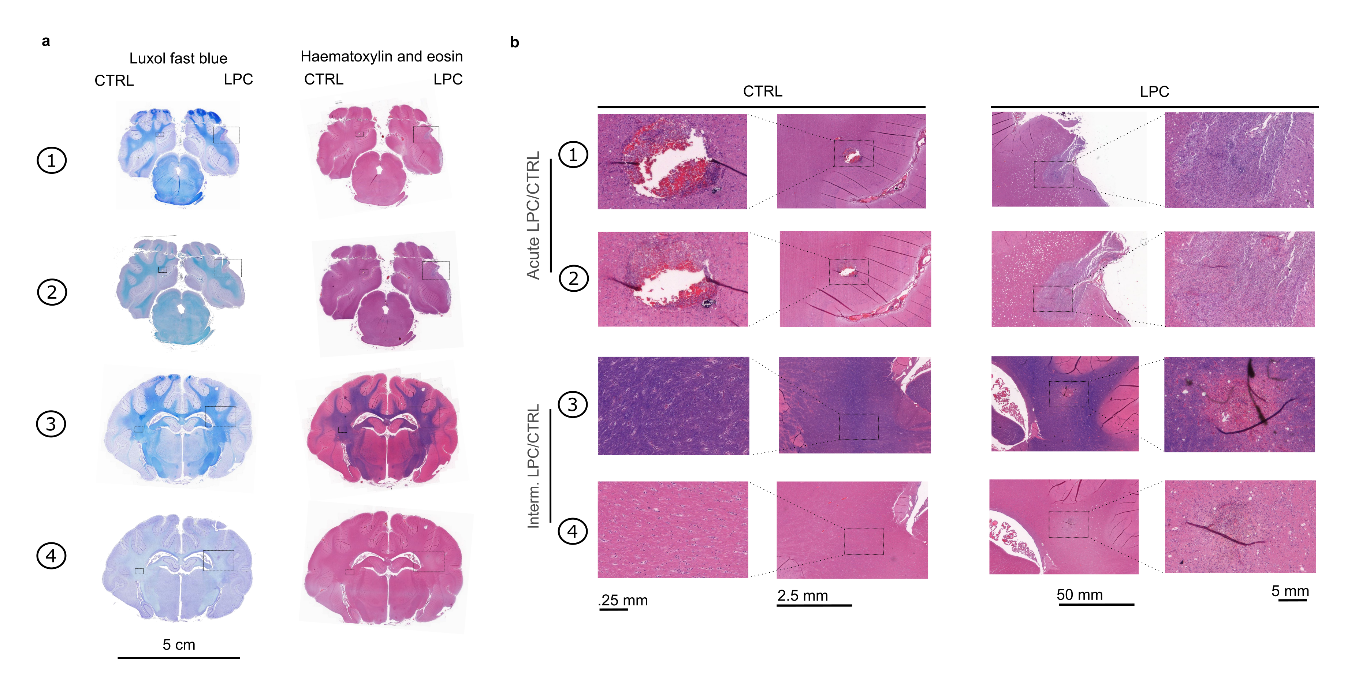

**Supplementary Figure 3.** *Histological section examples***.** a) Example of histopathological sections stained with Luxol fast blue (1^st^ column) and haematoxylin & eosin (2^nd^ column). Digits on the left correspond to the different planes from the 3D reconstruction in Fig. 4 as well as in b). The dotted rectangles represent the areas from which the higher magnifications on the right were taken. b) High magnification photomicrographs corresponding to the areas marked in a) (note different scales for presentation purposes, because LPC lesions are of an order of magnitude more extensive than CTRL).

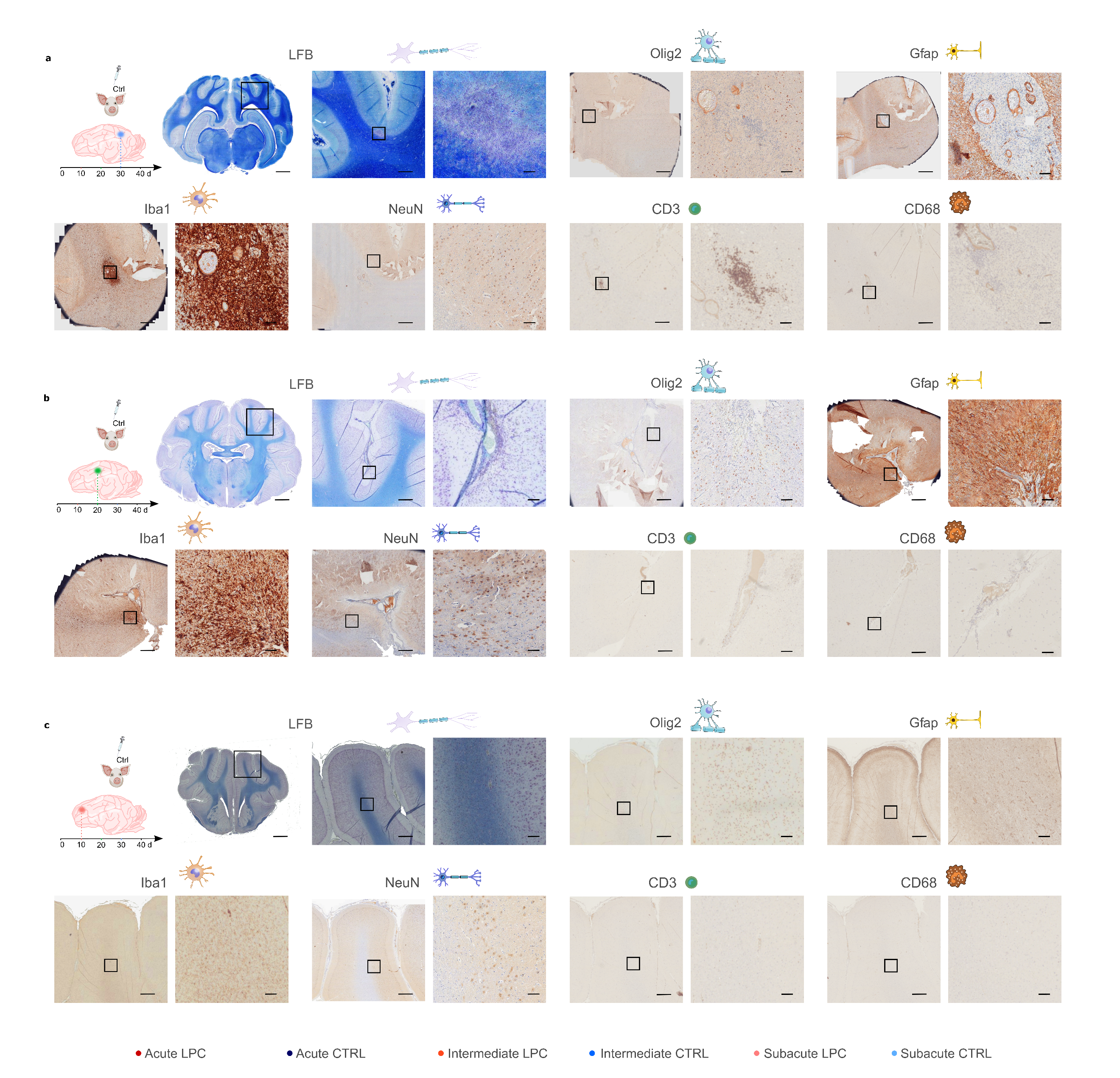

**Supplementary Figure 4.** *Immunohistochemical characterization of Ctrl lesions across stages.* a) Acute stage (*aCTRL*), b) Intermediate (*iCTRL*), c) Subacute (*sCTRL*) Common denominators of a)-c): From left to right: Schematic corresponding to Fig. 1 of the lesion stage; LFB (Luxol fast blue) overview of entire coronal slice (scale bar = 5 mm, black square delineates ROI magnified on the right and in each low magnification inset of the following stainings). Cell-specific marker stainings containing paired low-magnification (15x, left, scale bar = 200 µm, black square delineates ROI magnified on the right) and high magnification insets (right, 60x, scale bar = 50 µm) for following cellular markers: Olig2 (oligodendrocytes), Iba1 (microglia), NeuN (neuronal somata), CD3 (lymphocytes), CD68 (monocytes/macrophages).

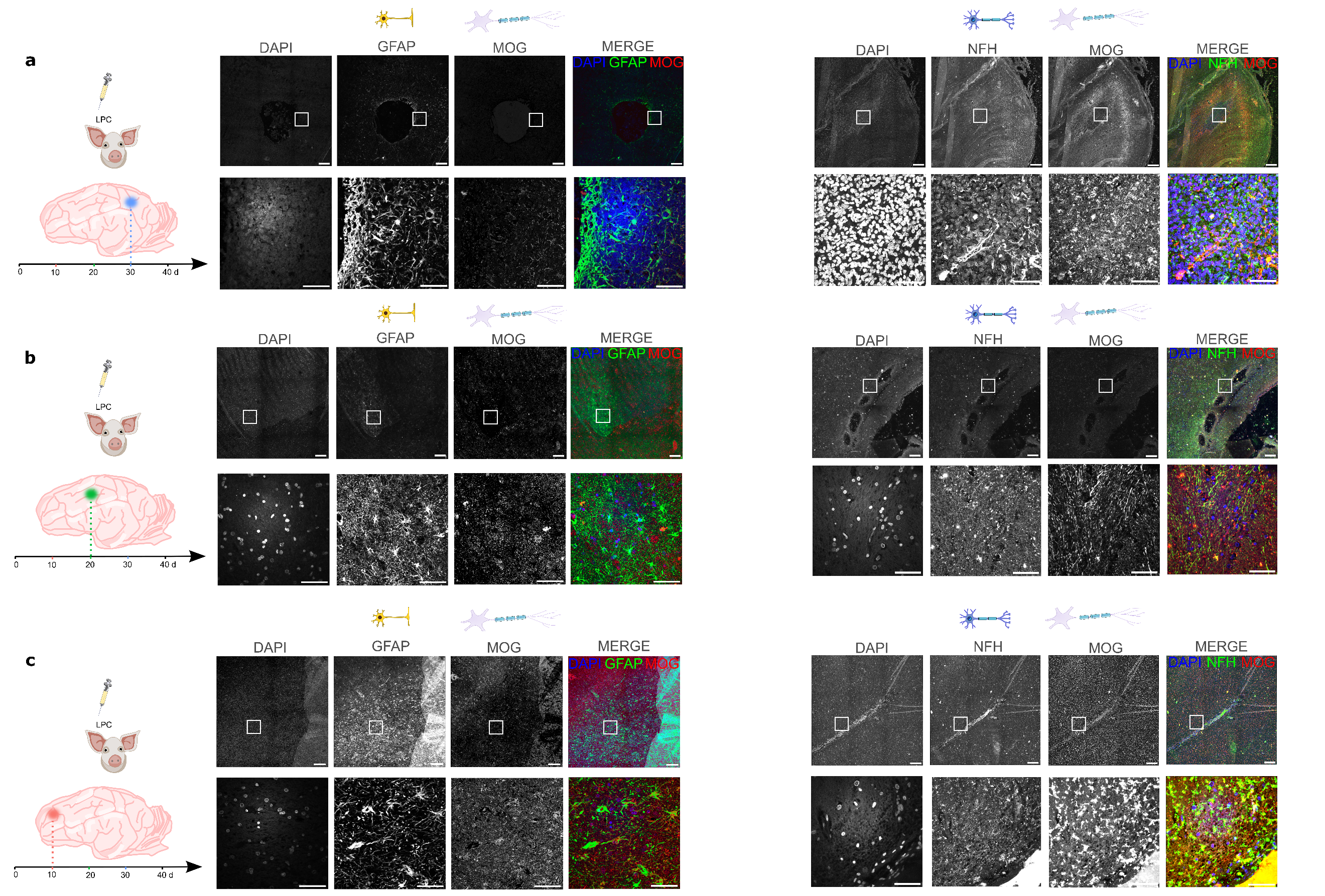

**Supplementary Figure 5.** *Immunofluorescence characterization of LPC lesions across stages.* a) Acute stage (*aLPC*), b) Intermediate (*iLPC*), c) Subacute (s*LPC*) Common denominators of a)-c): From left to right: Schematic corresponding to Fig. 1 of the lesion stage; DAPI (4′,6-diamidino-2-phenylindole, nuclear), GFAP (glial fibrillary acidic protein, astrocytic) and MOG (myelin oligodendrocyte glycoprotein, oligodendrocytic) triple stain (left half) and DAPI, NFH (neurofilament heavy chain, neuronal) and MOG triple stain (right half): low magnification – top row, 15x, scale bar = 200 µm, white box delineates ROI magnified in row below), high magnification – bottom row, 60x, scale bar = 50 µm.

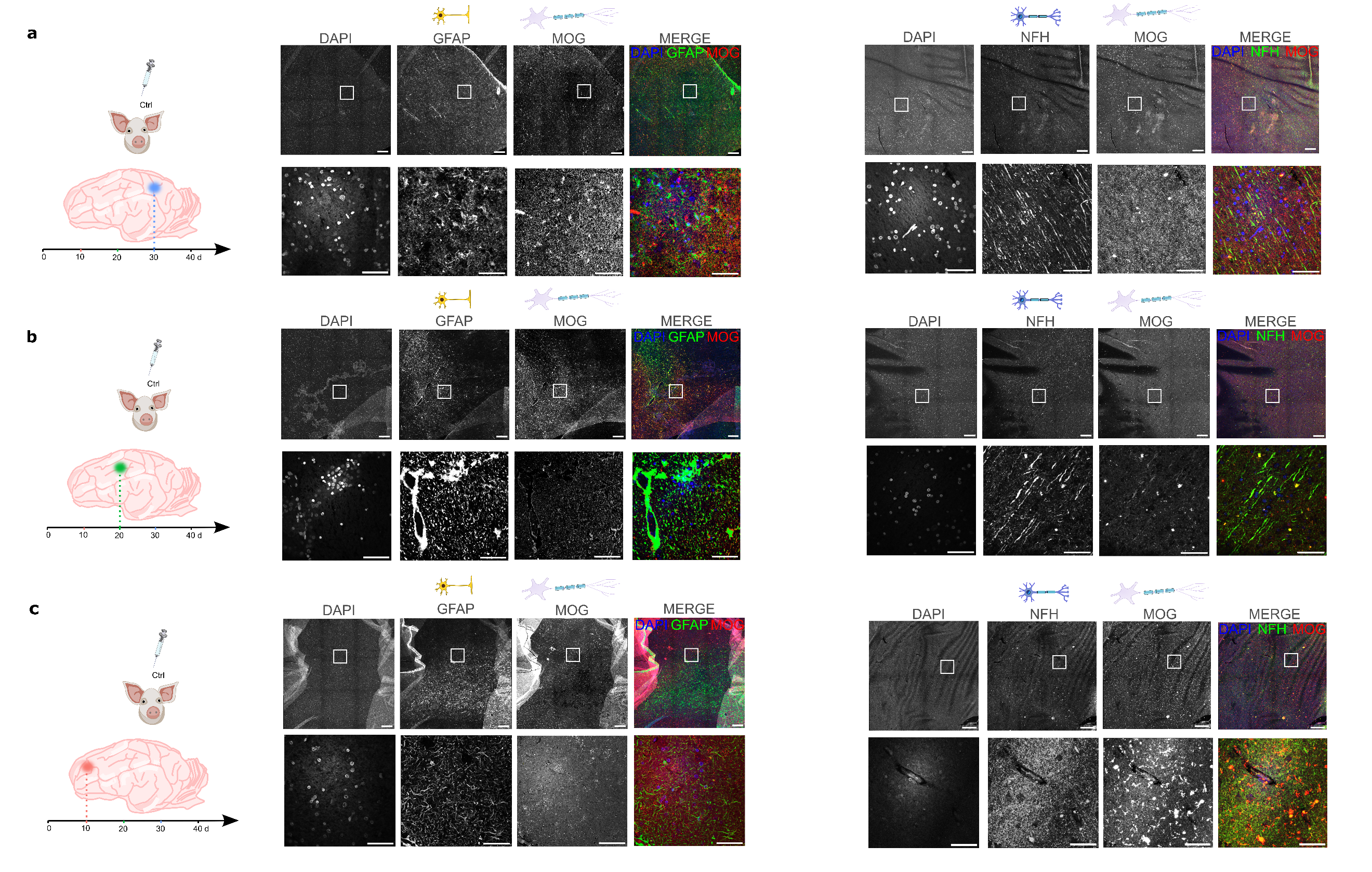

**Supplementary Figure 6.** *Immunofluorescence characterization of CTRL lesions across stages.* a) Acute stage (*aCTRL*), b) Intermediate (*iCTRL*), c) Subacute (*sCTRL*) Common denominators of a)-c): From left to right: Schematic corresponding to Fig. 1 of the lesion stage; DAPI (4′,6-diamidino-2-phenylindole, nuclear), GFAP (glial fibrillary acidic protein, astrocytic) and MOG (myelin oligodendrocyte glycoprotein, oligodendrocytic) triple stain (left half) and DAPI, NFH (neurofilament heavy chain, neuronal) and MOG triple stain (right half): low magnification – top row, 15x, scale bar = 200 µm, white box delineates ROI magnified in row below), high magnification – bottom row, 60x, scale bar = 50 µm.

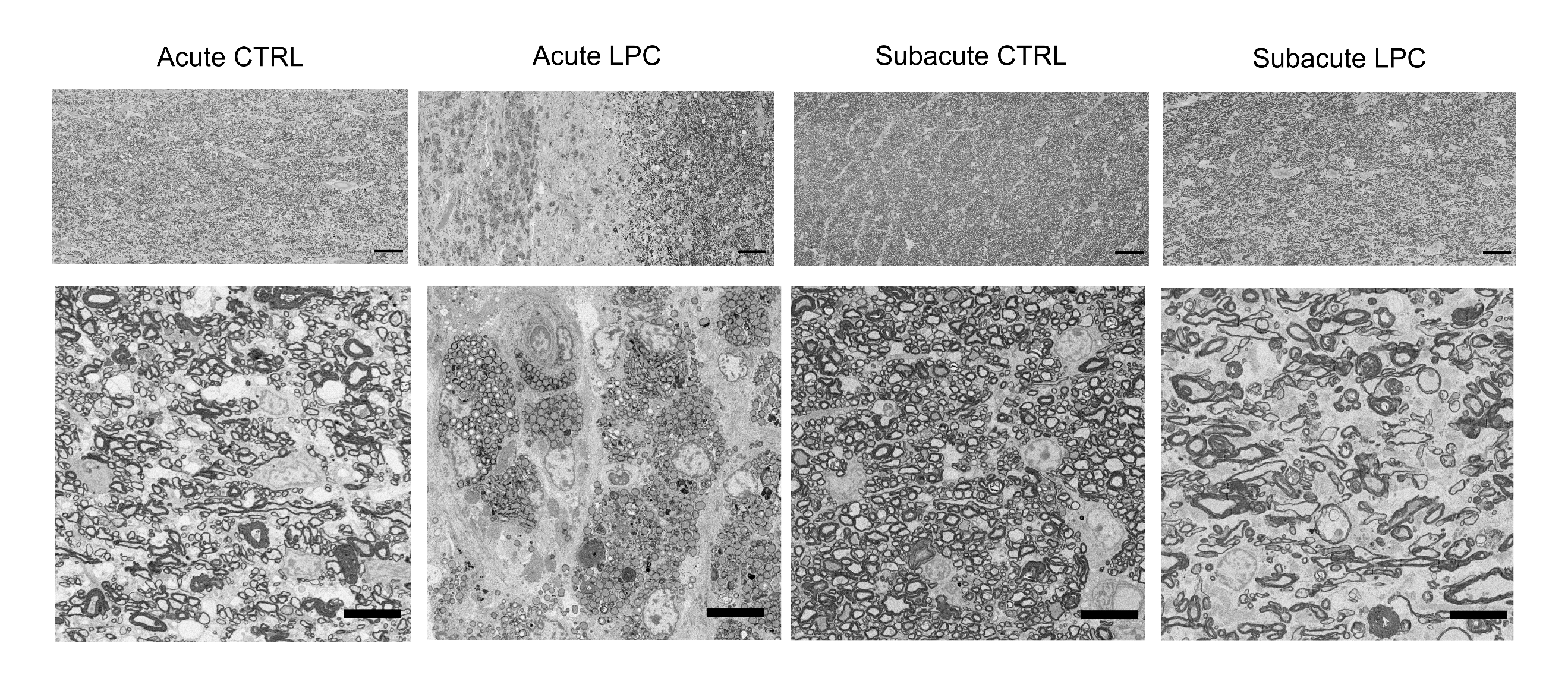

**Supplementary Figure 7.** *MiniSWINE* SEM-micrographs from *aCTRL, aLPC, sCTRL, sLPC* groups. Boxed areas in overview images (top) are shown at higher magnification (bottom). Scale bars 50 µm (top), 10 µm (bottom).
